# Oncodevelopmental plasticity of the skeleton in myeloid neoplasms

**DOI:** 10.64898/2026.03.19.712939

**Authors:** Salim Atakhanov, Ian Ghezzi, Hector Tejeda Mora, Lucas Greven, Marta Rizk, Lina Schmidt, Katrin Götz, Lina Merg, Valeria Solozobova, Adam Benabid, Pia Wanner, Niklas Lutterbach, Aleksandar V. Kargaliev, Mathias Schäfersküpper, Alexandru Florea, Juliette E. Pearce, Susanne Schmitz, Carmen Schalla, Paul Wanek, Rogerio B. Craveiro, Chloé Radermacher, Chiara Stüdle, Thomas Lehmann, Marek Weiler, Marcelo A.S. de Toledo, Steffen Koschmieder, Jitske Jansen, Francis Ayuk, Nicolaus Kröger, Felix M. Mottaghy, Daniel Truhn, Fabian Kiessling, Helene F.E. Gleitz, Tata Nageswara Rao, Michael Wolf, Carolin V. Schneider, Rafael Kramann, Anna Bock, Martina Crysandt, Michael Milsom, Rebekka K. Schneider

## Abstract

Myelofibrosis in patients with myeloproliferative neoplasms (MPNs) is traditionally characterized by bone marrow fibrosis and osteosclerosis, with *de novo* bone formation commonly attributed to impaired osteoclast-mediated resorption. Here, we challenge this paradigm by demonstrating that a solitary clonal driver mutation simultaneously induces pathological bone formation and resorption, with osteosclerosis acting to conceal localized and active bone destruction rather than inhibiting it. Through population analysis; clinical imaging; patient-derived multi-tissue sequencing; murine models and organ-on-a-chip systems, we demonstrate that spatial and ontogeny-dependent remodeling in mesoderm- and neural crest–derived bones is mechanistically interconnected via a previously unidentified osteochondral stromal injury program. Neural crest–derived stromal cells suppress osteogenic programs and undergo injury-induced lineage plasticity with ectopic chondrogenesis, mirroring pathological remodeling in mesoderm-derived growth plate regions. This shared injury response promotes osteoclastogenesis and is mediated by a conserved Thrombospondin 1+ (THBS1+) stromal population that links fibrotic remodeling to bone loss. Combined pharmacological inhibition of THBS1 and JAK signaling reduces myeloproliferation, halts fibrosis progression, and restores two developmentally distinct bones, establishing THBS1 as a unifying therapeutic target in myelofibrosis.

## Introduction

Clonal hematopoiesis of indeterminate potential (CHIP) provides a conceptual and biological continuum that links age-related premalignant states with overt myeloid disease. Among these lesions, JAK2V617F is uniquely pleiotropic: it arises in hematopoietic stem cells, promotes expansion in CHIP, and can manifest clinically as essential thrombocythemia (ET), polycythemia vera (PV), or primary myelofibrosis (PMF), while also serving as a common antecedent lesion of secondary acute myeloid leukemia^1^. Across this spectrum, JAK2V617F-mutant hematopoiesis confers a myeloid differentiation bias, chronic inflammatory milieu, and niche remodeling capacity, yet the full range of its effects on non-hematopoietic tissues remains incompletely understood.

One of the most neglected dimensions of JAK2V617F-driven CHIP is its impact on the skeleton. While myelofibrosis is well known for its profound osteosclerosis, observed in up to 70% of patients^2^, epidemiological data paradoxically reveal increased osteoporosis and fracture risk across the broader MPN spectrum^3,4,5^. Notably, osteoporotic fractures are preferentially associated with a fibrotic (and thus osteosclerotic) phenotype and JAK2-mutated disease^6^. Myeloid malignancies such as MDS and CMML also have a clear skeletal impact, including bone loss, fractures, and inflammatory joint disease^7^. Evidence from clonal hematopoiesis suggests that even premalignant clones can actively drive bone destruction through inflammatory mechanisms^8^. Thus, the same clonal driver mutation present across CHIP, MPN and myelofibrosis (MF) seems capable of producing opposing skeletal outcomes, such that bone loss and new formation may vary in severity across distinct anatomical sites and may even coexist within the same bone. However, current models of MPN pathophysiology do not explain how mutant hematopoiesis induces diverse skeletal phenotypes, and the role of osteochondral differentiation remains largely unexplored. We propose a paradigm-shifting hypothesis: that a single malignant clone can drive two seemingly opposing skeletal and stromal phenotypes - osteosclerosis and osteoporosis. This challenges the prevailing view that these outcomes are mutually exclusive or strictly disease stage dependent. Instead, we posit that dynamic bone-remodeling trajectories depend on the mutant clone-stromal crosstalk, potentially arising from a shared underlying stromal mechanism for divergent bone outcomes.

## Results

### Preferential periodontal bone involvement across ontogenetically distinct skeletal sites in MPN patients

To determine whether ontogenetically distinct bones undergo differential remodeling in MPNs, we performed a comprehensive whole-body analysis of ^18^F-labelled 2-fluoro-2-deoxy-D-glucose (FDG) positron emission tomography/computed tomography (PET/CT) scans from patients with myelofibrosis (PMF n=5, post-ET MF n=2) and age-matched controls (n=8);(Fig. 1a,b; Table S1a). Quantification of periodontal bone mineral density (BMD) showed a pronounced increase in the maxilla but only subtle changes in the mandible (Fig. 1c), revealing previously unrecognized craniofacial site-specific remodeling and underscoring the importance of analyzing the two jawbones separately.

**Fig. 1:**
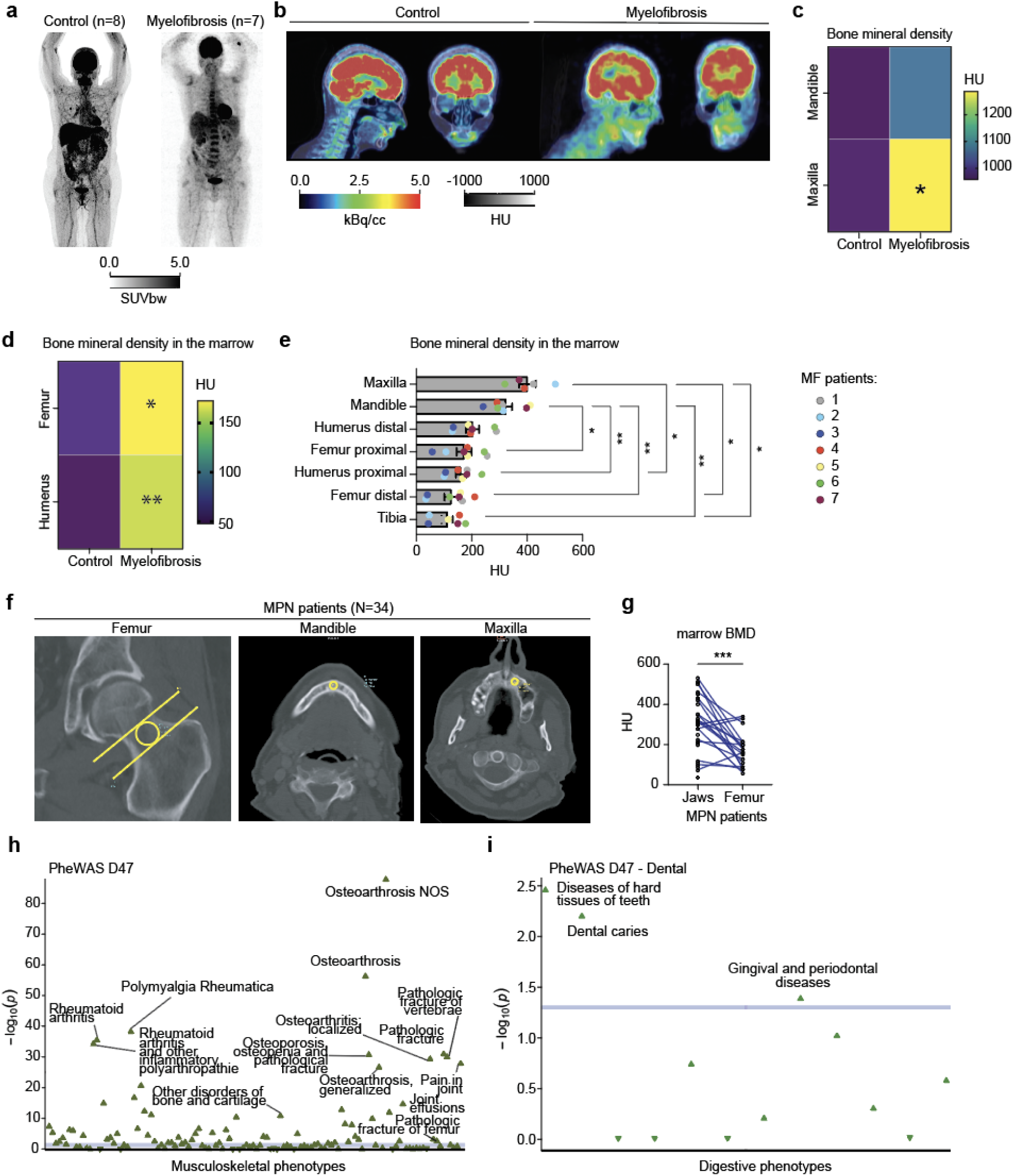
Preferential periodontal bone involvement across ontogenetically distinct skeletal sites in MPN patients. **a-b,** representative 18F-FDG PET/CT images of control (n=8) and myelofibrosis (n=7) patients. **c,** Quantification of bone mineral density of the maxillary and mandibular jaws in control and MF patients. HU=Hounsfield unit. Two-way ANOVA followed by Sidak’s multiple-comparisons test. **d,** Quantification of bone mineral density in the marrow of long bones (femur and humerus). HU=Hounsfield unit. Two-way ANOVA followed by Sidak’s multiple-comparisons test. **e,** Quantification of bone mineral density in the marrow of different skeletal sites of the same MF patients. Color code represents the parameters from each patient. HU=Hounsfield unit. One-way-ANOVA with post hoc Tukey’s was used. **f-g,** Representative images of retrospective radiologic CT analysis and quantification of bone mineral density in the marrow of the jawbones versus femur of the same MPN (n=34) patients. Statistical analysis was performed using paired two-tailed Student’s *t* test. **h, i,** Phenome-wide association study from UK Biobank focusing on musculoskeletal **(h)** and digestive phenotypes **(i)** of MPN individuals (n=2,548; ICD-10 D47).

Long-bone pathology was systematically evaluated in the femur and humerus. Both sites exhibited significantly increased marrow-associated BMD in MF, confirming a severe osteosclerotic phenotype in mesoderm-derived bones (Fig. 1d). To capture intra-patient heterogeneity, we extended our analysis to paired skeletal sites using whole-body PET/CT and retrospective clinical CT scans from a larger MPN cohort (n=34). Neural crest–derived bones (maxilla and mandible) consistently showed the highest marrow-associated BMD, whereas mesoderm-derived bones, particularly the tibia, displayed substantially lower values (Fig. 1e-g; Table S1b).

Assessment of marrow metabolic activity using FDG uptake revealed no global difference between MF patients and controls (Fig. S1a). Notably, the maxilla uniquely exhibited a significant reduction in glucose uptake. Given the well-established inverse relationship between marrow metabolic activity and fibrosis severity in MF^9,10^, this finding indicates pronounced bone remodeling within the periodontal niche.

While clinical imaging allows robust quantification of site-specific osteosclerosis and metabolic activity, its resolution limits the detailed analysis of trabecular microarchitecture and dynamic bone resorption. Notably, although osteoclast numbers are reportedly increased in MF, accumulating evidence suggests that their function is altered rather than absent, highlighting a potentially targetable imbalance in bone remodeling^11,12^. However, recent reports of malignancy-driven gingivitis and periodontitis in myeloid neoplasia challenge this assumption by demonstrating active osteolytic remodeling^13^. To overcome imaging limitations and systematically assess bone-destructive phenotypes at the population level, we leveraged UK Biobank (UKBB) data (∼500,000 participants) and identified 2,548 individuals with MPN (ICD-10 D47), enabling a phenome-wide association study (PheWAS). Across all significantly associated disease categories, osteoarthrosis emerged among the most strongly enriched musculoskeletal phenotypes in individuals with MPN (Fig. S1b). Restricting the analysis to musculoskeletal traits revealed significant associations with a broad spectrum of bone-destructive conditions, including rheumatoid arthritis, osteoarthritis, pathological fracture of the vertebrae and femur, osteoporosis, osteopenia, joint pain, and disorders of bone and cartilage (Fig. 1h). These findings at the population level indicate that MPN is associated with excessive bone formation but also with pervasive skeletal degeneration and fragility. Given the striking periodontal remodeling observed in imaging analyses, we next examined dental phenotypes. Disorders of tooth hard tissues, dental caries, and gingival and periodontal diseases were all significantly enriched in individuals with MPN (Fig. 1i).

Together, these findings demonstrate that MPN engages both osteosclerotic and bone-destructive programs in an ontogeny- and site-dependent manner and identify periodontal bone as a previously unrecognized skeletal niche with disproportionately severe disease involvement.

### Intra-patient single-cell mapping of ontogenetically distinct skeletal niches uncovers divergent cellular programs in myelofibrosis

Imaging and population-level analyses revealed marked differences in the severity of bone remodeling in MF, identifying periodontal bone as a uniquely affected skeletal niche. To define the cellular basis of these site-specific phenotypes, we performed scRNA sequencing of matched iliac crest bone marrow biopsies and extracted teeth from JAK2V617F-mutant MF patients (n=3) undergoing pre-transplant evaluation prior to allogeneic hematopoietic stem cell transplantation (Fig. 2a; Table S1c). These clinically annotated, paired samples obtained through routine dental screening and fibrosis grading enabled direct interrogation of hematopoietic–stromal interactions across ontogenetically distinct skeletal compartments. MF bone marrow was compared with age-matched orthopedic controls (n=3), while periodontal tissues were benchmarked against non-malignant periodontitis (n=3) and healthy reference samples (n=1) to delineate MPN-specific remodeling.

**Fig. 2:**
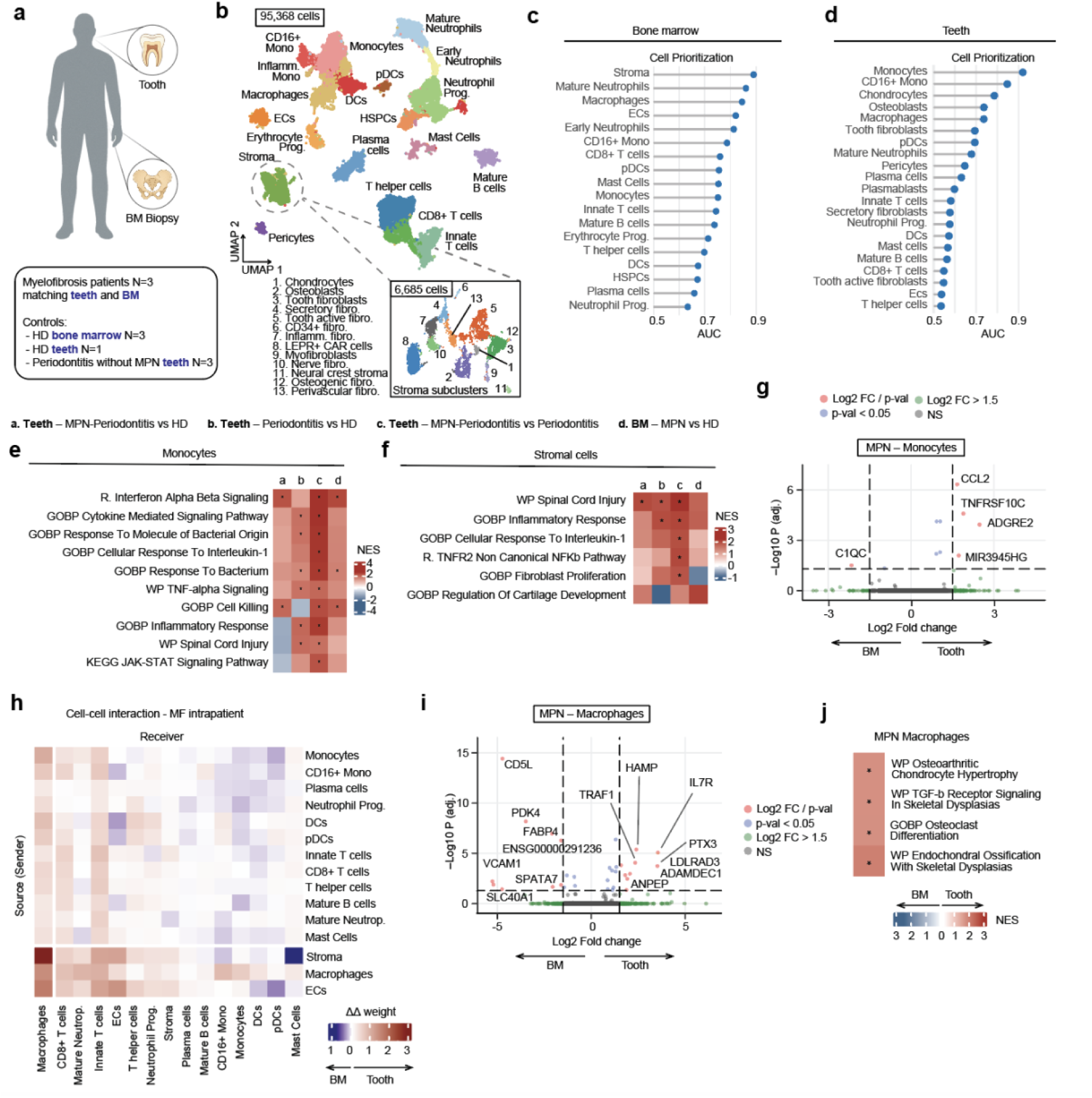
Intra-patient single-cell mapping of ontogenetically distinct skeletal niches uncovers cellular programs driving pathologic bone loss in MF. **a,** Schematic overview of patient cohorts, biopsy sites, and experimental design for intra-patient single-cell RNA sequencing of paired bone marrow (BM) and dental tissues from myelofibrosis (MF) individuals, including healthy donor (HD) controls and periodontitis-only controls where indicated. **b,** UMAP representation of the integrated single-cell transcriptomic dataset comprising 95,368 cells derived from paired BM and tooth biopsies. Inset shows refined clustering of 6,685 stromal cells resolved into 13 transcriptionally distinct subclusters. **c-d,** Cell prioritization analysis across all annotated cellular populations in BM **(c)** and teeth **(d)**, identifying the most perturbed cell types associated with MF-driven pathology in comparison with healthy controls. **e-f,** Pathway enrichment analysis of differentially expressed genes in monocytes **(e)** and stromal cells **(f)**, comparing conditions as follows: (a) MPN-associated periodontitis versus HD in teeth, (b) periodontitis without MPN versus HD in teeth, (c) MPN-associated periodontitis versus periodontitis without MPN in teeth, and (d) MPN versus HD in BM. Enrichment was performed using Reactome (R.), Gene Ontology (GO), WikiPathways (WP), KEGG, BioCarta, and PID databases. **g,** Volcano plot of differentially expressed genes (DEG) of monocytes from the intra-patient analysis comparing the expression of genes in the teeth versus the BM of the same MF individuals. **h,** Inference of intercellular communication networks in paired dental (red) and BM (blue) samples from the same MF individuals. **i,** Volcano plot of DEGs of macrophages from the intra-patient analysis comparing the expression of genes in the teeth versus the BM of the same MF individuals. **j,** Pathway enrichment analysis for macrophages comparing the teeth (red) versus the BM (blue) of the same MF individuals.

We profiled 95,368 single cells from sort-enriched bone marrow and dental samples across all conditions (Fig. 2a-b; Fig. S2a, Table S2, see Methods part). Unsupervised clustering identified 20 major cell populations encompassing hematopoietic progenitors, myeloid and lymphoid lineages, and non-hematopoietic stromal niche cells. Sub-clustering of 6,685 stromal cells resolved distinct fibroblast, osteogenic, chondrogenic, perivascular, neural, and CXCL12-abundant reticular (CAR) populations.

To identify cell types most responsive to MPN-associated injury, we applied Augur^14^, a machine-learning framework that quantifies transcriptional separability between conditions. Across both bone marrow and teeth, myeloid and stromal compartments consistently ranked among the most transcriptionally perturbed cell types in MPN (Fig. 2c, d). In dental samples, monocytes, CD16⁺ monocytes, and macrophages showed the strongest hematopoietic perturbations, whereas chondrocytes, osteoblasts, and tooth fibroblasts were the most affected stromal populations.

Pathway enrichment analysis revealed a pronounced inflammatory activation of monocytes in MPN-Periodontitis compared with periodontitis alone, marked by interferon, interleukin-1, TNF, and JAK–STAT signaling, alongside broader immune activation programs (Fig. 2e). Stromal cells from the same samples showed coordinated induction of inflammatory, fibroblast proliferation, and tissue injury pathways, consistent with a robust fibro-inflammatory state (Fig. 2f). Notably, cartilage development pathways were selectively enriched in MPN conditions in both teeth and bone marrow, indicating a shared osteochondral injury response across distinct skeletal niches. Consistent with these findings, monocytes from MPN-associated periodontitis upregulated pro-inflammatory genes including *ANKRD22, GBP5, OSM, CD274*, and *PIM1*, demonstrating that MPN amplifies local inflammatory programs beyond those observed in periodontitis alone (Fig. S2b-e).

We next asked whether MPN monocytes adopt distinct transcriptional states across skeletal environments. We performed differential expression analysis of monocytes isolated from pelvic bone marrow and teeth from the same MF patients, providing a direct comparison of clonal hematopoietic cells across ontogenetically distinct skeletal niches. Overall, monocytes from bone marrow and teeth showed highly similar transcriptional profiles, indicating that circulating MPN monocytes retain a conserved core identity across tissues (Fig. 2g). Despite this similarity, site-specific gene programs emerged. Bone marrow monocytes preferentially expressed complement-related genes, including *C1QC*, consistent with a role for complement activation in myelofibrosis. In contrast, periodontal monocytes upregulated genes linked to inflammation, tissue infiltration, and periodontitis progression (*CCL2, TNFRSF10C, MIR3945HG*), together with markers of monocyte-to-macrophage differentiation such as *ADGRE2*.

We applied CellChat to analyze cell–cell communication in bone marrow and periodontal tissues from the same individuals. Macrophages showed markedly increased incoming and outgoing signaling in the periodontium compared with bone marrow, with the strongest interactions occurring between macrophages and periodontal stromal cells (Fig. 2h). Tooth - resident macrophages expressed higher levels of genes linked to tissue infiltration and inflammation, consistent with an activated migratory state (Fig. 2i). Pathway analysis further revealed enrichment of bone-destructive programs, including osteoclast differentiation and endochondral ossification (Fig. 2j). Consistent with this, an osteoclast-like macrophage subpopulation was detected within the periodontal niche (Fig. S2f,g).

In summary, across distinct skeletal niches, MPN induces both shared and site-specific cellular programs, with the periodontal niche uniquely amplifying macrophage–stromal crosstalk and osteochondral activation, identifying the jaw as a previously unrecognized site of MPN-driven skeletal pathology.

### Myelofibrosis induces region-specific osteosclerosis and bone loss

To investigate if JAK2V617F mutant hematopoiesis alone is sufficient to cause skeletal pathology, we performed competitive bone marrow transplantation from inducible JAK2V617F or wild-type donors into wild-type recipient mice (Fig. 3a). Recipients of JAK2V617F marrow rapidly developed a PV–like phenotype, progressing to overt myelofibrosis by 20 weeks, characterized by splenomegaly, reduced bone marrow cellularity, and skewed hematopoietic differentiation toward myeloid progenitors (Fig. 3b,c; Fig. S3a-e). This shift included expansion of granulocyte–macrophage progenitors, a major source of osteoclast precursors, together with evidence of extramedullary hematopoiesis.

**Fig. 3:**
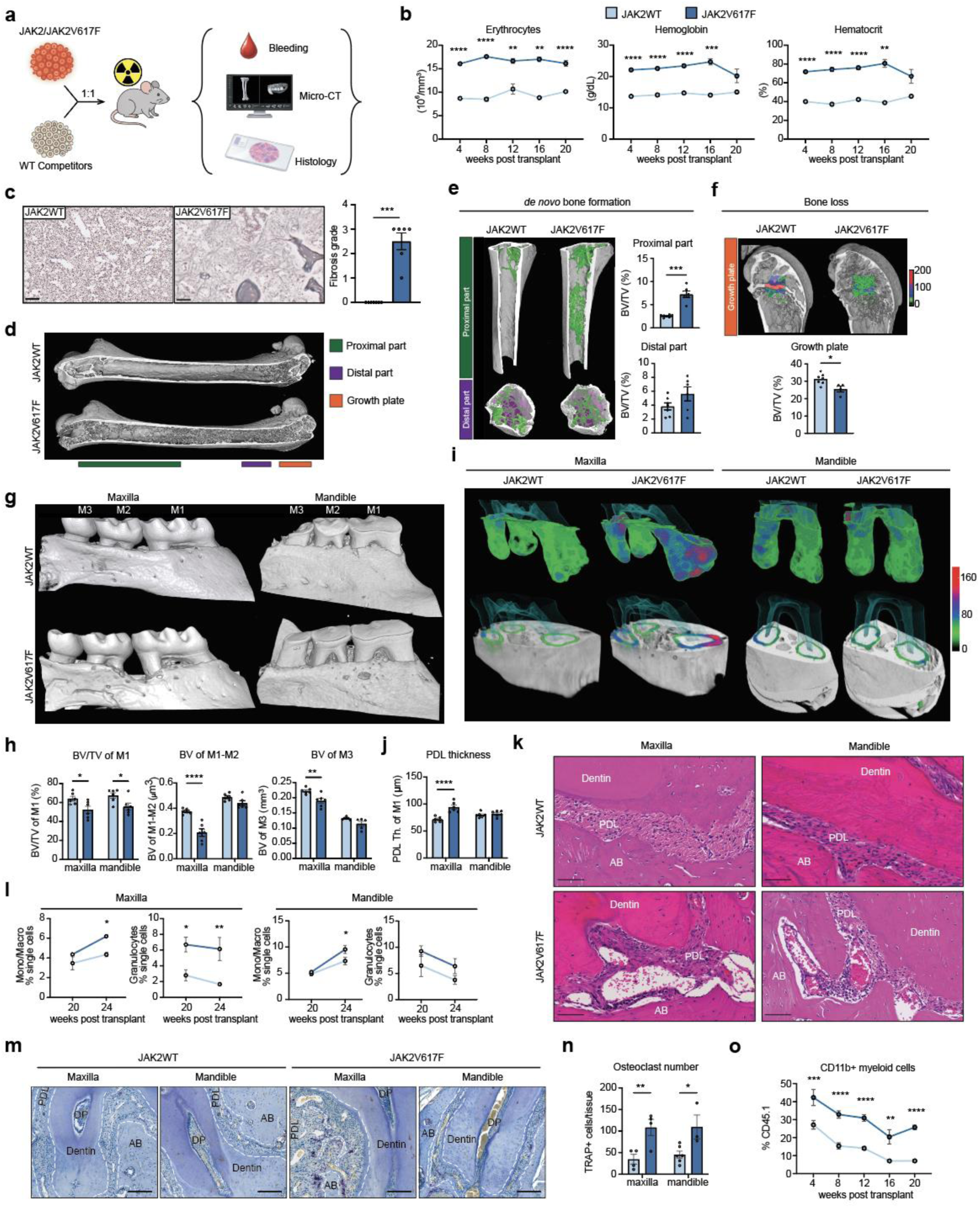
JAK2V617F-driven MF induces co-existing osteosclerosis and bone loss in ontogenetically distinct bones. **a,** Experimental design of the murine competitive transplantation assay. Whole bone marrow (BM) cells from JAK2WT or JAK2V617F donors were mixed 1:1 with wild-type (WT) competitor BM and transplanted into lethally irradiated WT recipient mice. Peripheral blood was collected every four weeks post-transplantation, and micro–computed tomography (µCT) and histological analyses were performed at 20 and 24 weeks post-transplantation. **b,** Longitudinal peripheral blood counts following transplantation (n = 6 mice per group). Data were analyzed by two-way ANOVA followed by Sidak’s multiple-comparisons test. **c,** Representative reticulin-stained BM sections (left) and quantification of reticulin fibrosis grade (right) in JAK2WT and JAK2V617F mice. Scale bar, 100 µm. **d-f,** Representative 3D reconstruction of micro-computed tomography (µCT) of distinct areas of interest within the murine femur (n=6 mice per group) and quantification of trabecular bone volume over total volume (BV/TV). **g-h,** Representative 3D µCT reconstructions of maxillary and mandibular bones from JAK2WT and JAK2V617F mice (n = 6) **(g)** and quantification of bone volume in defined regions of interest encompassing molar 1 (M1), the inter-molar region between molars 1 and 2 (M1–M2), and molar 3 (M3) **(h)** at 24 weeks post transplantation. **i, j,** Representative 3D images of periodontal ligament (PDL) thickness (Th.) beneath molar 1 of maxilla and mandible in both JAK2WT and JAK2V617F conditions **(i)** and its quantification **(j)** at 24 weeks post transplantation. **k,** Representative hematoxylin and eosin (H&E)–stained sections of the PDL region in maxillary and mandibular tissues from JAK2WT and JAK2V617F mice. Scale bar, 100 µm. **l,** Flow cytometric quantification of myeloid populations in maxillary and mandibular tissues over time, including monocytes/macrophages (CD11b+Gr1-) and granulocytes (CD11b+Gr1+), in JAK2WT and JAK2V617F mice (n = 6). Data were analyzed by two-way ANOVA followed by Sidak’s multiple-comparisons test. **m-n,** Representative tartrate-resistant acid phosphatase (TRAP) staining of periodontal tissues **(m)** and quantification of TRAP+ osteoclasts per tissue area **(n)** in JAK2WT and JAK2V617F mice at 24 weeks post transplantation. Scale bar, 50µm. PDL: periodontal ligament, DP: dental pulp, AB: alveolar bone. Statistical analysis was performed using two-way ANOVA with Sidak’s correction. **o,** Flow cytometric analysis of circulating mutant myeloid cells (CD45.1+CD45.2–CD11b+) in peripheral blood over time. Data were analyzed by two-way ANOVA followed by Sidak’s multiple-comparisons test.

To resolve how these hematopoietic alterations translate into bone pathology, we performed high-resolution micro–computed tomography (μCT) of mesoderm-derived long bones (femur) and neural crest–derived craniofacial bones (maxilla and mandible). The femur was subdivided into anatomically distinct proximal, distal and growth plate regions, enabling spatial resolution of bone remodeling patterns that are obscured by whole-bone analyses (Fig. 3d). μCT analysis revealed pronounced region-specific differences in bone remodeling within the same bone (femur). JAK2V617F mice developed pronounced osteosclerosis selectively in the proximal femur, whereas the distal region remained largely unaffected (Fig. 3e; Fig. S3f,g). Although bone volume was increased in the proximal femur, the trabeculae were thinner, indicating that bone resorption remains active even in osteosclerotic marrow. At the same time, the number of trabeculae was markedly increased, indicative of excessive new bone formation, and resulting in a dense but structurally immature trabecular network composed of many thin trabeculae in the proximal femur.

In contrast, the growth plate showed marked bone loss, with thinner and more widely spaced trabeculae and a modest shortening of the femur (Fig. 3f; Supplementary Fig. 3h,i). These findings suggest that bone regions closely linked to hematopoietic and stromal activity are selectively affected by MPN, whereas the mechanically specialized cortical bone is largely spared. Thus, bone gain and bone loss coexist within the same bone but are restricted to distinct functional compartments.

To determine whether these mechanisms extend to periodontal homeostasis, we analyzed jawbones at 20 and 24 weeks post-transplantation. At 20 weeks, mutant mice already exhibited early signs of an impaired periodontal bone remodeling, including increased osteoclast numbers, tendency in reduced bone volume around the first molar, thinner trabeculae and increased trabecular separation, indicative of an early resorptive phenotype (Fig. S3j-l). By 24 weeks, this progressed to severe periodontal bone loss affecting the entire maxilla with the subtle effect on the mandible (Fig. 3g,h), corroborating the clinical images of disproportional maxillary involvement in MF patients (Fig. 1c, e) and epidemiological association between MPN and periodontal disease (Fig. 1i). Accordingly, the connective tissue between the tooth and the alveolar bone, called the periodontal ligament (PDL), was significantly expanded beneath maxillary molar 1, while mandibular molar 1 exhibited bone loss without PDL expansion (Fig. 3l,j). Histological analysis revealed profound neo-angiogenesis within the PDL of both jaws, accompanied by disruption of fibroblast alignment between dentin and alveolar bone (Fig. 3k), a dental pathology predicted to destabilize the tooth and culminate in tooth loss.

Flow cytometric profiling of jawbones revealed severe myeloid infiltration, with increased monocyte frequencies correlating with elevated osteoclast abundance in both maxilla and mandible (Fig. 3l-n). Mutant CD45.1⁺CD45.2⁻CD11b⁺ osteoclast precursors were significantly increased in the peripheral blood (Fig. 3o).

We propose that the heightened vulnerability of the maxilla reflects its higher porosity and vascularization, enabling infiltration of (most likely circulating) osteoclast precursors and selective expansion of the maxillary PDL as a compensatory response to bone loss. These findings define a spatially segregated remodeling program in MPN in which pro-fibrotic (TGFb driven) programs in the marrow drive osteosclerosis, while mutant myeloid cells induce niche-specific osteoclastogenesis and periodontal bone loss. This uncoupling of bone formation and resorption provides a unifying mechanism for the coexistence of osteosclerosis, osteoporosis, and periodontitis in MPN, and, consistent with recent DNMT3A/CHIP studies^8^, supports the concept that clonal myeloid disease alone is sufficient to drive periodontal pathology.

### MPN drives ectopic chondrogenic fate of neural crest-derived stromal cells

To elucidate the mechanisms of JAK2V617F-driven periodontal remodeling, we performed scRNA sequencing on sorted cells from maxillae and mandibles of JAK2V617F and control mice (Fig. 4a). We profiled 15,790 cells spanning the major cellular compartments of the periodontium, generating a comprehensive single-cell atlas of murine maxillary and mandibular tissues. Unsupervised clustering identified 11 major cell populations comprising 45 subclusters, including neural, endothelial, epithelial, stromal, and hematopoietic lineages (Fig. 4b; Fig. S4a-c).

**Fig. 4:**
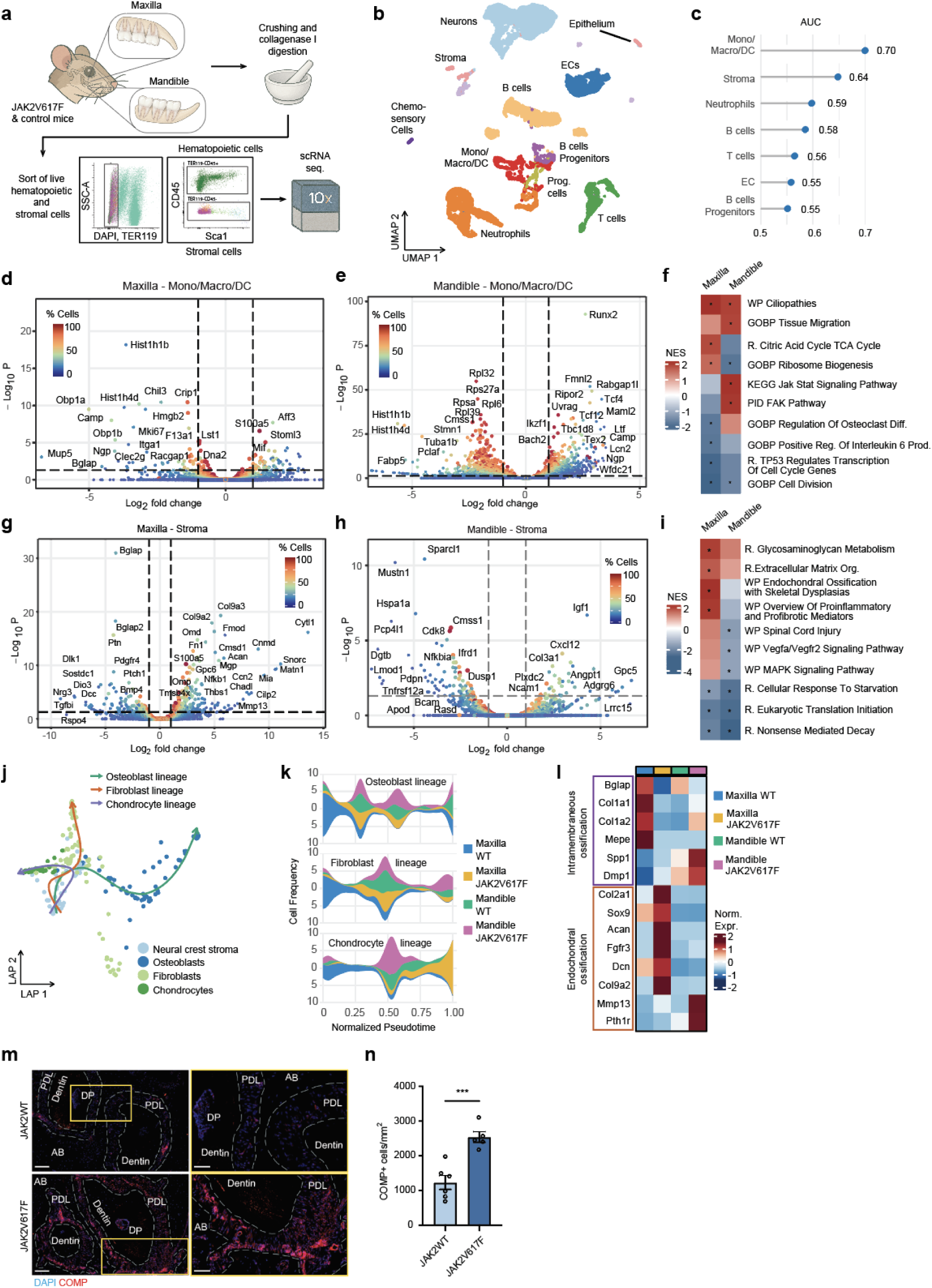
MPN causes the lineage plasticity of neural crest-derived stromal cells driving ectopic chondrogenesis and bone loss. **a,** Schematic overview of isolation and purification of hematopoietic and stromal cells from maxillary and mandibular bones of JAK2WT and JAK2V617F conditions for single cell RNA sequencing. **b,** UMAP representation of the integrated single-cell transcriptomic dataset from two jawbones in both conditions, comprising 15,790 cells resolved into 11 transcriptionally distinct subclusters. **c,** Cell prioritization analysis across broadly annotated cellular populations in both jawbones together, identifying the most perturbed cell types in JAK2V617F jaws in comparison with JAK2WT controls. AUC: area under curve. **d-e,** Volcano plot of differentially expressed genes (DEG) of mono/macro/DC population in maxilla **(d)** and mandible **(e)** comparing the expression of genes between JAK2V617F versus JAK2WT conditions. **f,** Pathway enrichment analysis for mono/macro/DC comparing JAK2V617F versus JAK2WT in maxilla and mandible. Pathway analysis was performed using Reactome (R.), Gene Ontology (GO), WikiPathways (WP), KEGG, BioCarta, and PID databases. **g-h,** Volcano plot of differentially expressed genes (DEG) of stromal population in maxilla **(g)** and mandible **(h)** comparing the expression of genes between JAK2V617F versus JAK2WT conditions. **i,** Pathway enrichment analysis for stromal cells comparing JAK2V617F versus JAK2WT in maxilla and mandible. Pathway analysis was performed using Reactome (R.), Gene Ontology (GO), WikiPathways (WP), KEGG, BioCarta, and PID databases. **j,** Trajectory inference of jawbone stromal subclusters using Slingshot, with inferred lineage paths overlaid onto dimensionality reduction plots. **k,** Visualization of condition-specific dynamics of stromal lineages using stream plots, with pseudotime values normalized across conditions. **l,** Distinct ossification-associated gene expression profiles of stromal cells across conditions in both maxilla and mandible. **m-n,** Representative immunofluorescence images of cartilage oligomeric matrix protein (COMP)–stained maxillary bones **(m)** and quantification of COMP+ cells per tissue area **(n)** in both conditions. PDL: periodontal ligament, DP: dental pulp. Scale bars, 100 µm (left) and 50 µm (right).

We applied Augur to rank cell states most perturbed by MPN-driven periodontal remodelling. Mono/macro/DC populations and stromal cells emerged as the two most responsive compartments, consistent with our patient-derived data (Fig. 2d). Differentially expressed gene (DEG) analysis of mono/macro/DC population revealed pronounced site-specific transcriptional reprogramming. In the maxilla, mono/macro/DCs upregulated osteoclastogenic genes (*Stoml3, MIF*), while downregulating osteogenic (*Bglap, Chil3, Crip1*) and cell-cycle–associated genes (*Mki67*), consistent with a resorptive, non-proliferative state (Fig. 4d). In contrast, mandibular mono/macro/DCs showed broad repression of ribosomal genes (*Rpl32, Rps27a, Rpsa, Rpl6, Rpl39*) together with induction of inflammatory, lineage-priming, and fate-determining regulators (*Tcf4, Tcf12, Lcn2, Ngp, Maml2*) (Fig. 4e). Pathway enrichment further revealed increased signatures related to tissue migration, alongside suppression of osteoclast differentiation, inflammatory and cell division pathways (Fig. 4f).

Periodontal stromal cells confirmed their neural crest origin and ontogenetic distinction from mesoderm-derived axial and appendicular bones (Fig. S4d), underscoring fundamental developmental differences of the craniofacial bones. Differential gene expression analysis revealed marked regional divergence in stromal responses. In the maxilla, stromal cells showed pronounced downregulation of osteogenic programs (*Bglap*, *Bglap2*, *Bmp4*, *Ptch1*, *Ptn* and *Tgfbi*) accompanied by upregulation of fibro-inflammatory (*Nfkb1*, *Fmod*, *Col9a2*, *Thbs1*), pro-osteoclastogenic signature (*Ccn2*, *Omd*), and chondrocyte-associated genes (*Acan*, *Cytl1*, *Cnmd*, *Chadl*, *Mia*, *Matn1*, *Snorc*, *Mmp13*) (Fig. 4g). In contrast, mandibular stromal cells downregulated inflammatory and stress-response pathways (*Nfkbia*, *Tnfrsf12a*, *Dusp1*, *Pdpn*), and stromal structural and musculoskeletal genes (*Sparcl1*, *Mustn1*). Instead, they showed enrichment of myeloid recruitment (*Cxcl12*) and angiogenic support programs (*Igf1*, *Angpt1*, *Adgrg6*) (Fig. 4h), consistent with histological and immunophenotyping analysis (Fig. 3k-n).

Pathway enrichment analysis corroborated these findings, revealing increased extracellular matrix organization, endochondral ossification with skeletal dysplasia signatures, alongside enrichment of pro-inflammatory and pro-fibrotic mediators, coupled with suppression of cellular stress adaptation pathways, including responses to starvation and nonsense-mediated decay (Fig. 4i). Concordantly, periodontal stromal cells displayed an elevated profibrotic module score without induction of bone formation programs, indicating a maladaptive stromal response driving periodontal bone loss (Figure S4e,f).

We next investigated the unexpected activation of chondrocyte-associated gene programs in maxillary stromal cells. As chondrocytes are absent from periodontal maxillary bone, which forms exclusively by intramembranous ossification, this aberrant signature was striking. Its consistent presence in both patient and murine datasets prompted a focused analysis of stromal cell fate reprogramming.

Trajectory and pseudotime analyses resolved three neural crest–derived stromal lineages: osteoblastic, fibroblastic, and chondrocytic (Fig. 4j). Notably, the chondrocyte trajectory emerged exclusively under JAK2V617F-driven hematopoietic stress (Fig. 4k), identifying ectopic chondrogenesis as a pathological stromal response to mutant hematopoiesis.

To define the mesenchymal differentiation programs underlying this fate switch, we curated gene signatures associated with intramembranous versus endochondral ossification. Interestingly, JAK2V617F-exposed maxillary stromal cells adopted an aberrant chondrocyte transcriptional state, activating ectopic endochondral ossification machinery within the periodontal niche and downregulating the intramembranous ossification signature (Fig. 4l). Aberrant chondrogenesis in the maxilla was further validated at the protein level by robust expression of cartilage oligomeric matrix protein (COMP) in the PDL in partially spindle-shaped, partially rounded cells with abundant cytoplasm (Fig. 4m,n). The number of COMP+ cells was significantly increased in the JAK2V617F condition.

Given the femoral growth plate bone loss (Fig. 3f) and our single-cell evidence for expansion of proliferative pre-chondrocytes in murine myelofibrosis^15^, we asked whether chondrocytes in mesoderm-derived bones undergo similar pathological reprogramming as in the maxilla. To address this, we leveraged our previously published scRNAseq datasets of sorted TdTomato⁺ stromal cells isolated from long bones of three independent Cre-reporter lines (Pdgfrb;tdTomato, Gli1;tdTomato and Grem1;tdTomato)^15^. These datasets span BM fibrosis induced by thrombopoietin (ThPO) overexpression and control conditions. We therefore focused on the long bone-derived chondrocyte compartment to test whether fibrotic marrow remodeling induces shared pathological programs across ontogenetically distinct bones. Differential gene expression analysis of mesoderm-derived chondrocytes revealed marked suppression of osteogenic and bone-supportive genes (*Bglap, Col1a1, Clec11a, Sparc*), accompanied by downregulation of the hematopoietic niche factor *Cxcl12* (Fig. S4g). Concomitantly, these cells upregulated fibro-inflammatory and immunomodulatory programs (*Acta2, Lcn2, Tsc22d3, Cytl1, Txnip*), consistent with an injury-associated chondrocyte state. Notably, this transcriptional profile closely mirrored that of ectopic neural crest–derived chondrocytes in the maxilla, including shared repression of osteogenesis (*Bglap*) and promotion of osteoclastogenesis (*Cytl1*)^16^, indicating convergence on a common osteochondral injury program despite distinct developmental origins in long bones and jaw. Pathway enrichment analysis further substantiated this phenotype, revealing suppression of skeletal system development, ossification, and cartilage programs involved in endochondral bone morphogenesis, alongside enrichment of apoptotic pathways, a hallmark of degenerative chondrogenesis observed in osteoarthritis^17^ (Fig. S4h). Together, these findings align with μCT-detected growth plate defects and mechanistically link aberrant chondrocyte states to bone-destructive remodeling in MF.

Collectively, we identify a conserved osteochondral injury program across ontogenetically distinct bones in MF, driven by convergent pathological reprogramming of neural crest–derived stromal cells and mesoderm-derived chondrocytes. This shared chondrocyte-centered response serves as a unifying mechanism for MPN-associated skeletal pathology, encompassing both arthritis and periodontitis.

### A conserved THBS1⁺ stromal cell population links myelofibrosis to osteochondral remodeling

To understand how cellular interactions instruct pathological differentiation of neural crest stromal cells in MPN, we next interrogated intercellular communication within the periodontal niche. We applied CellChat to systematically infer cell–cell interaction networks in both the maxilla and mandible (Fig. 5a; Fig. S5a). In addition to the neurovascular inflammatory pathways (Fig. S4i,j), neuronal and endothelial compartments showed pronounced communication hubs in JAK2V617F-driven dental pathology, suggesting a highly activated neurovascular communication axis during MPN-associated periodontal bone loss.

**Fig. 5:**
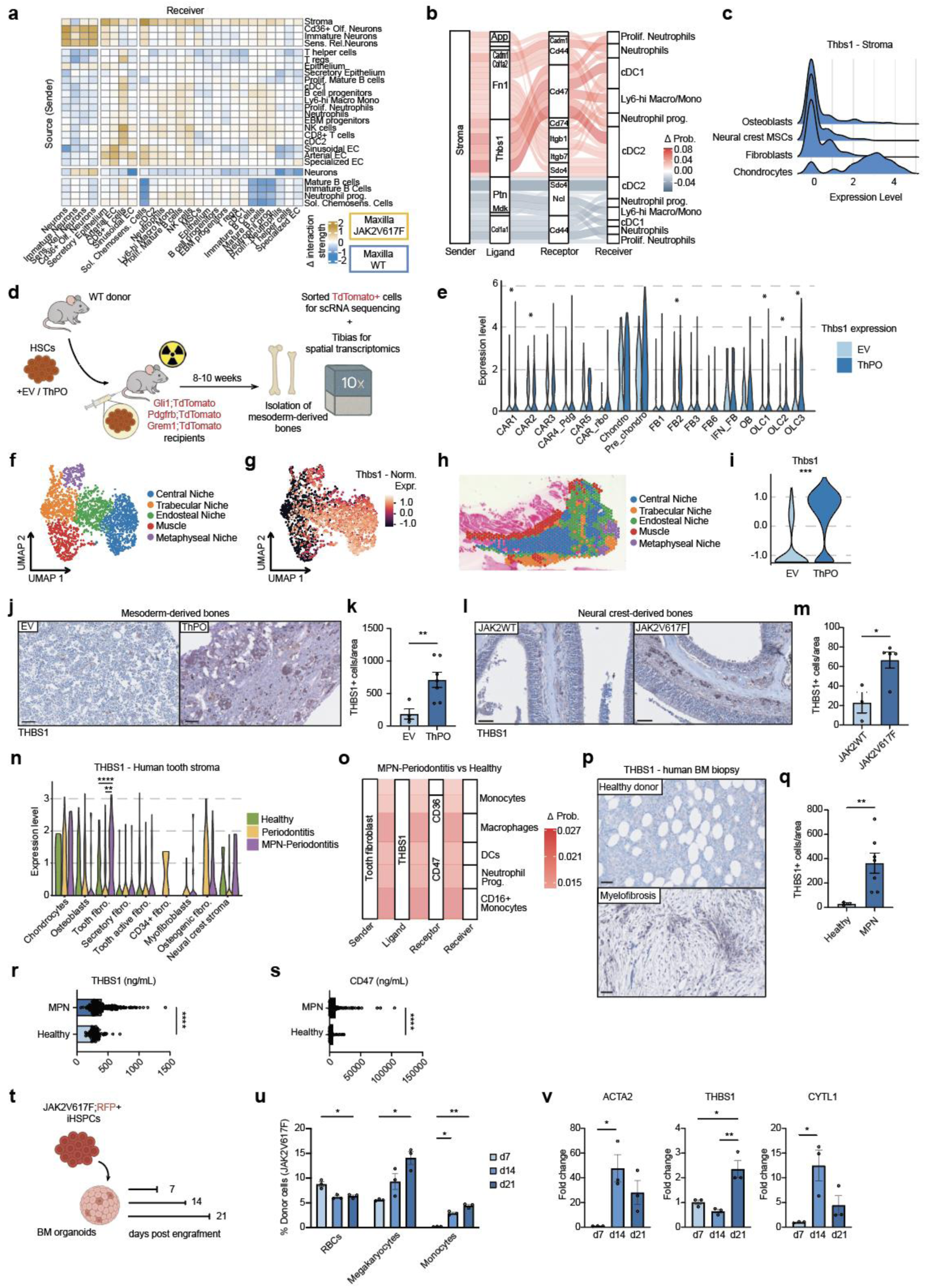
A conserved THBS1⁺ stromal population mediates injury responses across ontogenetically distinct bones in myelofibrosis. **a,** Intercellular communication networks in paired maxillary jawbones comparing JAK2V617F versus JAK2WT conditions. **b,** Cell-cell communication depicting ligand (L)-receptor(R) interaction between maxillary stromal cells (sender) and myeloid cells (receiver). Interactions are represented based on the difference in delta probability of mean LR expression between JAK2V617F and JAK2WT. **c,** Thrombospondin 1 (Thbs1) expression across transcriptionally defined stromal subclusters in the maxilla. **d,** Schematic overview of the murine thrombopoietin (ThPO)–driven myelofibrosis model. c-Kit–enriched hematopoietic stem cells isolated from WT donors were lentivirally transduced with ThPO or empty vector (EV) and transplanted into bigenic CreER;tdTomato reporter recipients (Gli1;tdTomato, Pdgfrb;tdTomato and Grem1;tdTomato). Following tamoxifen induction and lethal irradiation, recipients received donor cells intravenously. TdTomato⁺ stromal cells were isolated from mesoderm-derived bones for single-cell RNA sequencing, and tibias from the same cohorts were used for Visium spatial transcriptomics. **e,** Thbs1 expression in 17 annotated stromal subclusters under fibrotic (ThPO) and control (EV) conditions in mesoderm-derived bones. **f-g,** UMAP representation of integrated Visium spatial transcriptomics data from ThPO and EV tibias (n = 3 mice per group), identifying five major spatial domains **(f)** and corresponding Thbs1 expression patterns **(g)**. **h,** Representative overlay image of H&E and spatial domains detected in tibias from Visium spatial transcriptomics, n=3 per group. **i,** Thbs1 expression from Visium spatial transcriptomics dataset in both fibrosis (ThPO) and control (EV) conditions. **j-k,** Representative images of Thbs1-stained tibias in fibrosis (ThPO) and control (EV) groups **(j)** and the quantification of Thbs1+ cells per tissue area **(k)**. Scale bars, 50µm. **l-m,** Representative images of Thbs1-stained maxillary bones in both JAK2 and JAK2V617F conditions **(l)** and the quantification of Thbs1+ cells per tissue area **(m)**. Scale bars, 50µm. **n,** Thbs1 expression across nine annotated periodontal stromal subclusters from human tooth biopsies of MPN-periodontitis (n = 3), periodontitis without MPN (n = 3), and healthy controls (n = 1). Stromal subclusters are ordered according to cell-state prioritization by Augur (from left to right). **o,** Cell-cell communication depicting ligand (L)-receptor(R) interactions between patients’ tooth fibroblast subpopulation (sender) and myeloid cells (receiver). Interactions are represented based on the difference in delta probability of mean LR expression between MPN-Periodontitis (N=3) and healthy control (N=1) patients. **p-q,** Representative images of Thbs1-stained BM biopsies from MPN (N=7) and healthy (N=3) patients **(p)** and the quantification of Thbs1+ cells per tissue area **(q)**. Scale bars, 50µm. **r-s,** ELISA of Thbs1 **(r)** and its interaction partner CD47 **(s)** in plasma of MPN (N=293) and healthy (N=74) individuals. **t,** Schematic overview of 3D in vitro experiment. JAK2V617F;RFP+ iPSCs were differentiated into CD34+ cells and engrafted into mature healthy iPSC-derived BM organoids. Organoids were harvested 7, 14 and 21 days post engraftment of exogenous cells. **u,** Flow cytometry analysis depicting the frequency of donor-derived (JAK2V617F;RFP+) red blood cells (RBCs), megakaryocytes and monocytes. Cells were isolated from BM organoids after 7, 14 and 21 days post engraftment of exogenous cells. Two-way-ANOVA with post hoc Tukey’s was used. **v,** qRT-PCR analysis from BM organoid-derived cells after 7, 14 and 21 days post engraftment of exogenous cells. One-way-ANOVA with post hoc Tukey’s was used.

Stromal cells in both the maxilla and mandible emerged as dominant signal-sending hubs within the periodontal niche, positioning them as central organizers of pathogenic intercellular communication during MPN-driven bone loss (Fig. 5a; Fig. S5a). This observation prompted us to examine whether stromal cells actively instruct myeloid behavior through specific ligand–receptor programs. Ligand–receptor interaction analysis uncovered a robust and selective stromal–myeloid signaling axis mediated by thrombospondin-1 (THBS1)–CD47 axis (Fig. 5b). *THBS1* expression was markedly upregulated in maxillary stromal cells during periodontal bone loss (Fig. 4g), consistent with our prior findings implicating *THBS1* in mutant megakaryocyte–stromal crosstalk in human myelofibrosis^18^. Stromal subcluster-resolved analysis revealed that *THBS1* expression was highest within the ectopic chondrocyte population (Fig. 5c), identifying these injury-induced stromal cells as a dominant signaling source within the periodontal niche. Pathway enrichment of THBS1-expressing chondrocytes highlighted programs associated with extracellular matrix organization, integrin-mediated cell–surface interactions, responses to elevated platelet cytosolic Ca²⁺, and metabolic disease pathways (Fig. S5b), consistent with a pro-fibrotic, maladaptive stromal state.

To define the regulatory circuitry underlying *THBS1* induction in MPN, we used decoupleR to infer transcription factor (TF) associations. This analysis revealed that RUNX3 and SMAD2, key effectors of TGF-β signaling, were positively correlated with THBS1 expression^19^, whereas YY1 was negatively associated (Figure S5c).

Given that chondrocytes from two ontogenetically distinct bones showed a common osteochondral injury program, we investigated whether stromal THBS1 induction is a conserved response to myelofibrosis across skeletal compartments. To address this, we again leveraged the previously published and above-mentioned scRNA-seq datasets from the ThPO overexpression and empty vector (EV) control models (Fig. 5d). Across 17 transcriptionally distinct stromal subclusters, *THBS1* expression was significantly elevated in myelofibrotic conditions, with strongest induction in CAR1/2 cells, fibroblast subset FB2, osteolineage clusters (OLC1-3) and particularly in chondrocyte and pre-chondrocyte populations (Fig. 5e), suggesting differentiation of neural crest-derived stromal cells into ectopic THBS1^high^ chondrocytes in response to periodontal injury.

Visium spatial transcriptomics on tibias of the same cohort highlighted five distinct spatial regions by unsupervised clustering: central, trabecular, endosteal, metaphyseal and muscle - associated (Fig. 5f, Fig. S5d). *THBS1* was highly enriched within the central, endosteal and metaphyseal regions (Fig. 5g). Spatial mapping onto hematoxylin–eosin–stained sections revealed that the THBS1-enriched endosteal domain localizes to the growth plate and metaphyseal regions and significantly increased in ThPO condition (Fig. 5h,i). Immunostaining confirmed robust THBS1 expression in both mesoderm-derived long bones and neural crest–derived jawbones in MPN (Fig. 5j-m). In long bones, THBS1 localized to the fibrotic interstitium and dysplastic megakaryocytes, whereas in the maxilla THBS1+ cells were enriched in subendosteal regions, supporting a convergent stromal response across ontogenetically distinct bones. In human periodontal tissue, our scRNA-seq revealed MPN-specific THBS1 induction in tooth fibroblasts, the third most perturbed stromal population (Fig. 2d), but not in periodontitis alone or controls (Fig. 5n). CellChat and ligand–receptor analyses identified stromal-myeloid cell interactions mediated by conserved THBS1–CD36 and THBS1–CD47 signaling axes as well as selective, MPN-specific stromal–osteoclast communication (Fig. 5o; Fig. S5e).

To determine whether *THBS1* expression extends beyond neural crest–derived bones in humans, we analyzed pelvic bone marrow biopsies from MPN patients. We performed THBS1 immunohistochemistry (IHC) staining on FFPE bone marrow sections from MPN patients and healthy donors. Healthy bone marrow displayed normal cellularity with preserved adipocyte content and minimal THBS1 staining. In contrast, myelofibrotic marrow exhibited profound hypocellularity, extensive fibrosis, and a significant accumulation of THBS1⁺ cells (Fig. 5p,q). We further measured circulating THBS1 and its interaction partner CD47 in peripheral blood plasma from MPN patients and healthy controls using ELISA. Both THBS1 and CD47 protein levels were significantly elevated in MPN plasma compared with controls (Fig. 5r,s), indicating both local and systemic activation of this pathway potentially mediated by circulating platelets and remnants of stromal cells^20^.

To define the temporal induction of *THBS1*, we modeled MPN evolution using human BM organoids engrafted with isogenic JAK2V617F or JAK2WT iHSPCs^21,22^. To recapitulate stromal–hematopoietic crosstalk relevant to periodontal disease, we isolated PDL fibroblasts from wisdom teeth of a healthy individual and established an immortalized human PDL fibroblast line with preserved osteogenic differentiation potential and stable proliferative capacity across passages (Fig. S5f,g). This allowed integration of BM organoids with tooth-associated stromal cells in a multi-organ context.

We next engrafted the PV-patient derived JAK2V617F iPSC line and its CRISPR-repaired isogenic control (JAK2WT) in organoids^23^. For fate tracing within the BM organoids, JAK2V617F iPSCs were lentivirally labeled with RFP and isogenic JAK2WT controls with GFP, followed by directed differentiation into CD34⁺ hematopoietic stem and progenitor cells (iHSPCs). JAK2V617F iHSPCs displayed enhanced proliferation and altered maturation dynamics compared with isogenic controls (Fig. S5h).

Fluorescently tagged and purified CD34⁺ iHSPCs were engrafted into fully mature BM organoids made with a newly generated healthy iPS cell line (Fig. S5i-k). To recreate systemic communication, BM organoids were co-cultured with PDL fibroblast spheroids on a microfluidic multi-organ-on-a-chip platform, with physical separation allowing interaction exclusively via circulating factors and migratory cells, thereby mimicking blood-mediated signaling *in vivo* (Fig. S5l). Early PV-like phenotypes showed no THBS1 induction in organoids or PDL fibroblast spheroids despite altered hematopoiesis (Fig. S5m-p). In contrast, prolonged mutant clone exposure at 14 and 21 days post engraftment resulted in progressive lineage skewing toward megakaryocytic and monocytic fates, and coincident upregulation of *ACTA2, THBS1*, and the pathologic chondrocyte-associated gene *CYTL1*, paralleling fibrotic remodeling (Fig. 5t-v). Thus, THBS1 represents a late-stage, niche-intrinsic response to chronic mutant hematopoietic stress rather than an early consequence of myeloproliferation.

### Pharmacologic inhibition of THBS1 reverses dual skeletal pathologies in myelofibrosis

Having identified THBS1 as a unifying stromal mediator linking osteosclerosis and osteoporosis across ontogenetically distinct bones, we tested whether pharmacological THBS1 inhibition could mitigate MPN-associated fibrosis and aberrant bone remodeling. We employed LSKL to pharmacologically inhibit THBS1 activity. LSKL is a small peptide, which mimics a sequence within the latency-associated peptide (LAP) of TGFβ and selectively antagonizes THBS1-mediated TGFβ activation without directly inhibiting TGFβ signaling^19,24^, consistent with enrichment of THBS1-associated extracellular matrix (ECM) and TGFβ programs (Fig. S5b,c).

We competitively engrafted JAK2V617F;RFP and isogenic JAK2WT;GFP iHSPCs into fully mature human bone marrow organoids at a 1:1 ratio, followed by treatment with LSKL or vehicle control (Fig. 6a). In vehicle-treated organoids, mutant cells rapidly developed a robust MPN phenotype characterized by expansion of progenitor populations, increased immature and mature erythrocytes, and accumulation of megakaryocytes and monocytes (Fig. 6b; Fig. S6a). In striking contrast, LSKL treatment selectively reduced the abundance of mutant JAK2V617F HSPCs and attenuated erythrocytosis, while sparing isogenic WT competitor cells. Transcriptomic profiling of treated organoids revealed coordinated suppression of key fibrotic and inflammatory programs, including *ACTA2, COL1A1, IL1β*, alongside suppression of *THBS1* and its interaction partner *CD47* (Fig. 6c). Notably, LSKL treatment also reduced expression of chondrogenic markers, including *ACAN* and *SOX9*, indicating inhibition of the pathological stromal reprogramming observed during disease progression.

**Fig. 6:**
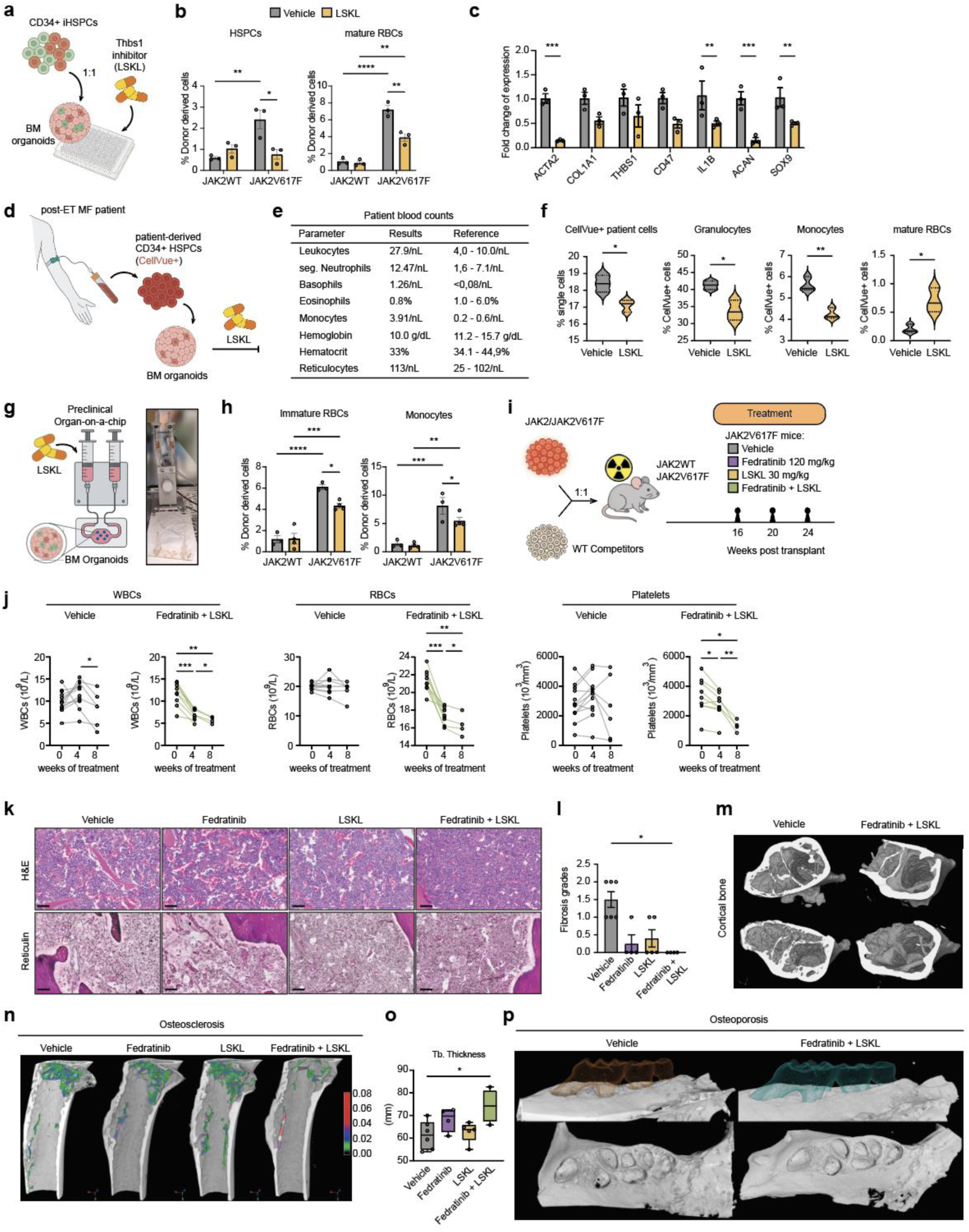
Pharmacological targeting of THBS1 synergizes with JAK inhibition to suppress myeloproliferation, halt fibrosis, and restore bone homeostasis. **a,** Experimental design of the *in vitro* BM organoid treatment assay. Mature human iPSC-derived bone marrow organoids were competitively engrafted with JAK2V617F;RFP⁺ and isogenic JAK2WT;GFP⁺ differentiated hematopoietic stem and progenitor cells (iHSPCs) at a 1:1 ratio and treated statically with the THBS1 antagonist LSKL or vehicle control for 7 days. **b,** Flow cytometric analysis of donor-derived cells from BM organoids treated with vehicle or LSKL, showing frequencies of HSPCs (CD45+CD34+) and erythroid cells (CD45-CD235a+) derived from mutant and WT competitors. Two-way-ANOVA with post hoc Tukey’s was used. **c,** qRT-PCR of BM-derived cells following LSKL or vehicle control treatment. Statistical analysis was performed using two-way ANOVA with Sidak’s correction. **d,** Schematic overview of treatment effect on patient-derived primary cells engrafted into BM organoids. Peripheral blood mononuclear cell-derived CD34+ fraction from a patient diagnosed with post-ET MF were labelled with CellVue dye and engrafted into mature iPSC-derived BM organoids. BM organoids with patient cells were treated statically with the THBS1 antagonist LSKL or vehicle control for 7 days. **e,** Clinical hematological parameters showing blood counts of a post-ET MF patient at the time of PBMC isolation. **f,** Flow cytometric analysis of CellVue-labeled patient-derived cells following vehicle or LSKL treatment. **g,** Schematic overview of the preclinical organ-on-a-chip platform enabling dynamic drug perfusion. BM organoids engrafted with JAK2V617F;RFP and JAK2WT;GFP CD34+ cells were cultured under continuous flow, with LSKLor vehicle delivered exclusively through the microfluidic circulation channel. **h,** Flow cytometric analysis of the preclinical organ-on-a-chip model with vehicle or LSKL treatment groups, showing frequencies of immature erythrocytes (CD45+CD235a+) and monocytes (CD45+CD14+) derived from mutant and WT competitors. Two-way-ANOVA with post hoc Tukey’s was used. **i,** Experimental design of in vivo therapeutic intervention. Lethally irradiated mice were competitively transplanted with JAK2V617F/JAK2WT and WT competitor bone marrow cells at a ratio 1:1. Blood counts were performed once in 4 weeks starting from 4 weeks post transplant. JAK2V617F mice started receiving the treatment at 16 weeks post-transplantation (time point 0) with vehicle (n=6), Fedratinib (n=4), LSKL (n=5), or combined Fedratinib + LSKL (n=4) until 20 and 24 weeks post transplant (4 and 8 weeks of treatment, respectively). **j,** Blood counts over time showing white blood cells (WBC), red blood cells (RBCs) and platelets. Each dot represents the value from one mouse. Two-way-ANOVA with post hoc Tukey’s was used. **k,** Representative images of H&E and reticulin staining on tibial sections from mice treated for 8 weeks. Scale bars, 50µm. **l,** Reticulin staining and grading of bone marrow fibrosis from mice treated for 8 weeks with vehicle (n=6), Fedratinib (n=4), LSKL (n=5), or combined Fedratinib + LSKL (n=4). Kruskal-Wallis H test was used. **m-n,** Reconstruction of 3D µCT images of femoral cortical (M) and trabecular (N) bones from mice treated for 8 weeks. **o,** Quantification of trabecular (Tb.) thickness from the proximal part of the femur. Mice treated for 8 weeks with vehicle (n=6), Fedratinib (n=4), LSKL (n=5), or combined Fedratinib + LSKL (n=4). One-way-ANOVA with Holm-Sidak’s correction was used. **p,** Reconstruction of 3D µCT images of maxillary bones from mice treated for 8 weeks.

To assess clinical relevance, primary cells from a patient diagnosed with post–essential thrombocythemia myelofibrosis (post-ET MF) were engrafted in organoids. The patient presented with marked leukocytosis characterized by neutrophilia and basophilia, a leukoerythroblastic blood film with 7% circulating blasts as well as severe hypochromic, microcytic anemia (Fig. 6d,e). Patient peripheral blood-derived CD34⁺ HSPCs were labeled with CellVue fluorescent dye, engrafted into mature BM organoids and treated *ex vivo* with either LSKL or vehicle control. Vehicle-treated cultures recapitulated the patient’s granulocytic bias and erythroid failure, whereas LSKL markedly reduced mutant myeloid expansion and partially restored erythroid differentiation (Fig. 6f; Fig. S6b,c), demonstrating efficacy in primary human disease.

To model physiological drug delivery, we implemented a perfusion-based organ-on-a-chip platform (Fig. 6g). In this system, mature BM organoids competitively engrafted with JAK2V617F;RFP and isogenic JAK2WT;GFP iHSPCs were placed in a dedicated tissue chamber, while LSKL or vehicle was administered exclusively through the perfusion set on the microfluidic unit, allowing the compound to circulate and penetrate the organoid in a diffusion-and flow-dependent manner. Consistent with static cultures, vehicle-treated organoids developed a pronounced MPN phenotype driven by the mutant clone, characterized by expansion of erythroid cells, monocytes, megakaryocytes and HSPCs (Fig. 6h, Fig. S6d). Strikingly, perfusion-based LSKL treatment significantly attenuated this phenotype. In addition to reducing erythroid expansion, LSKL robustly suppressed monocytic output, indicating that THBS1 inhibition effectively mitigates a PV phenotype with associated monocytosis under physiologically relevant delivery conditions. This highlights the organ-on-a-chip platform as a powerful preclinical tool for translational drug evaluation.

As bone marrow organoids lack a mature bone compartment, we next sought to evaluate the impact of THBS1 inhibition on fibrosis and bone remodeling *in vivo*. We hypothesized that simultaneous targeting of mutant hematopoietic cells and the fibrotic stromal niche, through combined JAK2 inhibition and THBS1 antagonism, could achieve superior therapeutic benefit.

Using a competitive transplantation model, in which secondary myelofibrosis develops between 16 and 20 weeks post-transplantation (Fig. 3b), we defined this interval as a therapeutic window for intervention. Lethally irradiated mice were reconstituted with a mixture of JAK2V617F/JAK2WT and wild-type competitor cells and monitored until 16 weeks post-transplantation (Fig. 6i). Mice were then treated with vehicle, the JAK2-selective inhibitor (fedratinib) alone, LSKL alone, or a combination of fedratinib and LSKL for either 4 weeks (until week 20 post transplant) or 8 weeks (until week 24 post transplant).

Prior to treatment initiation, all JAK2V617F mice exhibited pronounced myeloproliferation, with elevated white blood cell counts, platelets, erythrocytes, hemoglobin, and hematocrit (Fig. S6e). While monotherapies showed limited benefit, combined fedratinib + LSKL uniquely normalized blood counts, including sustained platelet correction (Fig. 6j, Fig. S6f). Histopathology revealed normalization of vascular architecture, reduced megakaryocyte clustering, and significant inhibition of fibrosis progression exclusively with combination therapy (Fig. 6k,l). Strikingly, micro–CT analysis demonstrated that combination therapy also rescued pathological bone remodeling, correcting coexisting osteosclerotic and osteoporotic lesions in the femur (Fig. 6m-o) and restoring alveolar bone integrity in the maxilla (Fig. 6p).

Collectively, these findings identify THBS1 as a unifying and disease-modifying therapeutic target across ontogenetically distinct skeletal compartments. Combined THBS1 antagonism and JAK inhibition not only restrain myeloproliferation and fibrotic progression but also restore bone homeostasis by simultaneously correcting stromal-driven osteosclerosis and mutant clone–driven osteoporosis, two opposing skeletal pathologies that co-exist within the same bone in myelofibrosis. Thus, our study uncovers THBS1 as a central node linking hematopoietic malignancy, osteochondral injury and aberrant bone remodeling, providing a therapeutic avenue to normalize BM fibrosis and skeletal pathology in MPN.

## Discussion

MPNs have traditionally been defined by bone marrow fibrosis and osteosclerosis, fostering the prevailing assumption that bone loss is largely absent in these disorders. Here, we challenge this paradigm by demonstrating that MPNs engage spatially segregated and ontogeny-dependent bone remodeling programs, in which osteosclerosis and bone loss not only coexist but are mechanistically linked through a previously unrecognized osteochondral injury response.

By integrating high resolution imaging, single cell sequencing approaches, murine and 3D *in vitro* models, we show that MPN induces both shared and niche-specific cellular programs across ontogenetically distinct skeletal compartments. In mesoderm-derived long bones, classical osteosclerosis within the marrow niche coexists with pronounced trabecular thinning and growth plate–associated bone loss, indicating that osteoclast-mediated resorption remains active despite apparent sclerosis. Building on our previous work^15^, we propose a niche-dependent model in which fibrotic injury redistributes osteolineage support away from the growth plate toward the diaphysis, promoting osteosclerosis while simultaneously unleashing localized osteoclast-driven bone destruction. This spatial uncoupling provides a mechanistic explanation for the long-standing underappreciation of bone loss in chronic MPNs: osteosclerosis does not replace bone destruction but instead masks ongoing, regionally confined skeletal degeneration.

We identify osteochondral injury as a unifying, bone-destructive program operating across developmentally distinct skeletal compartments. In neural crest–derived maxillary bone, which normally forms through intramembranous ossification, stromal cells display exceptional lineage plasticity in response to chronic mutant hematopoietic stress. They aberrantly adopt a metaphyseal-like, chondrogenic transcriptional state rather than sustaining osteogenesis. We propose that this chondrogenic shift occurs at the expense of osteoblast commitment, impairing local bone formation and creating a permissive microenvironment for excessive osteoclastogenesis. Notably, this mechanism mirrors the pathological remodeling at the growth plate, revealing a shared osteochondral injury response across distinct skeletal tissues. These findings extend prior observations that neural crest–derived stromal cells are highly plastic^25,26^ and activate chondrogenic programs in response to injury^27,28,29^, demonstrating that such lineage plasticity is pathologically hijacked in MPN to drive periodontal bone loss.

Clinical imaging in MPN patients and µCT analyses in mice demonstrate that the maxilla is disproportionately affected compared with the mandible. This differential susceptibility aligns with fundamental anatomical and developmental distinctions, as the maxilla forms a highly trabeculated intramembranous bone, whereas the mandible develops in association with Meckel’s cartilage and acquires a thicker cortical architecture, conferring distinct stromal remodeling properties.

Across both ontogenetically distinct bones, we identify THBS1⁺ stromal cells as a conserved effector population selectively enriched within osteolytic and fibrotic niches. These cells engage conserved signaling axes with myeloid populations, positioning THBS1 as a central mediator linking stromal injury responses, aberrant extracellular matrix remodeling, and osteoclast-driven bone loss in myelofibrosis.

Consistent with this role, THBS1 has been broadly implicated in inflammaging and myeloid bias of HSCs^30^, in shaping immunosuppressive microenvironments across malignancies and shown to be involved in TGFβ–SMAD signaling^31,32^. In solid tumors, THBS1 promotes T cell exhaustion through CD47 engagement, and its loss within the tumor microenvironment partially restores sensitivity to immune checkpoint blockade and cytotoxic therapies^33^. In myeloid malignancies, such as myelodysplastic neoplasms, stromal THBS1 contributes to an immunosuppressive microenvironment^34^.

Clinically approved drugs for MPNs, including JAK inhibitors, provide symptomatic relief but are not disease-modifying and have minimal impact on bone marrow fibrosis or pathological bone remodeling^12,18,35,36^. Consequently, there is a substantial unmet clinical need for therapies that can restore skeletal homeostasis while also halting fibrotic progression. Pharmacological inhibition of THBS1 has been successfully explored in multiple preclinical settings, including solid tumors^33^ and models of organ fibrosis^19^. Here, we demonstrate that THBS1 inhibition in MPN, in combination with clinically approved JAK inhibition, simultaneously attenuates myeloproliferation, halts fibrotic progression, and restores two opposing skeletal phenotypes. This underscores the disease-modifying potential of targeting stromal injury pathways.

In summary, we establish skeletal pathology in MPN as a spatially structured, injury-driven process governed by pathological osteochondral remodeling and stromal–myeloid crosstalk. By identifying THBS1 as a conserved, disease-modifying stromal effector operating across ontogenetically distinct bones, this work unifies fibrosis, aberrant skeletogenesis, and bone loss into a single mechanistic framework and defines a tractable therapeutic axis to restore marrow and skeletal homeostasis in myeloproliferative disease.

## Acknowledgments

The authors thank the involved team members in Aachen, Rotterdam and Heidelberg, IZKF Aachen in the Medical Faculty of RWTH Aachen, iPSC Facility at Erasmus MC and C.Thanisch (Ibidi GmbH) for experimental and technical assistance. The data used in this publication was managed using the research data management platform Coscine with storage space of the Research Data Storage (RDS) (DFG: INST222/1261-1) and DataStorage.nrw (DFG: INST222/1530-1) granted by the DFG and Ministry of Culture and Science of the State of North Rhine-Westphalia. R.K.S. is an Oncode Investigator and is supported by ERC grants (Rewind-MF ERC-CoG 101124542; deFIBER ERC-StG 757339 and PoC DeAlarmin) and a ZonMW VIDI grant. This work was in part supported by grants of the Deutsche Forschungsgemeinschaft (DFG) (German Research Foundation) to R.K. (KR 4073/9-1), R.K.S. (504777725; 417911533; 514007497), Mi.W. (504777725), S.K. (KO 2155/7-2; AOBJ 690056; project no. 417911533), F.K. (403224013; 331065168), M.M. (CRC873 INST 35/1899-1) and MPN Research Foundation to M.M. R.K.S received funding for the project from the program “Netzwerke 2021”, an initiative of the Ministry of Culture and Science of the State of North Rhine Westphalia (CANTAR network). R.K. and R.K.S. are members of the E:MED Consortia Fibromap and the consortium CureFib funded by the German Ministry of Education and Science (BMBF). TNR acknowledges the support from the Swiss National Science Foundation (320030-231993); Krebsforschung Schweiz (KFS-6308-02-2025-R); Wilhelm Sander-Stiftung (Förderantrags Nr. 2023.139.1). This research has been conducted using the UK Biobank Resource under Application Number 245437. UK biobank data was accessed by C.V.S. *Copyright © 2025, NHS England.* Re-used with the permission of the NHS England and/or UK Biobank. All rights reserved. This work uses data provided by patients and collected by the NHS as part of their care and support.

## Author contributions

S.A., I.G., H.T.M. designed, performed *in vivo*, *in vitro* experiments and biocomputational analyses. S.A. conceptualized the study, collected all data, wrote the original draft, designed the figures. L.G., J.J., R.B.C., C.R., M.A.S.T. performed *in vitro* experiments. M.R., Ma.W., F.K., Mi.W acquired and analyzed µCT data. H.T.M., L.S., J.E.P. analyzed scRNAseq and spatial transcriptomics data. L.M., V.S., K.G., Ad.B., Pi.W., N.L., H.F.E.G performed *in vivo* experiments. S.S., C.S., Pa.W., S.B. assisted with immunophenotyping and histological experiments. A.V.K., D.T. performed retrospective, radiologic CT analysis of patients. M.S., A.F., F.M.M, F.A., N.K. acquired and analysed clinical PET/CT images. Ch.S., T.L., T.N.R. provided FFPE BM biopsies and performed ELISA. Mi.W., An.B., M.C., S.K. identified patients, annotated the clinical samples and acquired biopsies. C.V.S. performed population-level study. Mi.W., S.K., F.K., T.N.R., R.K., M.M., R.K.S. acquired funding. M.M., R.K.S. conceptualized and supervised the study, wrote and edited the text.

## Methods

### Human samples

For single cell studies and formalin fixed paraffin-embedded (FFPE) tissues from bone marrow biopsies and teeth only excess material was used. In accordance with the approval from the ethics committee of the Medical Faculty of the RWTH Aachen University (EK24-409, EK094/16, EK173/06, EK300/13, EK127-12) all patient material was de-identified at inclusion. Unprocessed bone marrow biopsies and extracted teeth were obtained approximately 2h after successful sample acquisition via the Department of Hematology, Oncology, Hemostaseology and Stem Cell Transplantation, the Department of Oral and Maxillofacial Surgery as well as the Department of Orthodontics in the Faculty of Medicine of the RWTH Aachen University. More detailed information on patient characteristics can be found in Table S1c.

Whole-body PET/CT images of the myelofibrosis patients were provided from the University Medical Center Hamburg-Eppendorf with the ethics approval number 2022-100964-BO-ff. Control samples were retrospectively selected and provided by the Department of Nuclear Medicine at the RWTH Aachen University with the ethics approval number EK24-409. More detailed information on patient characteristics can be found in Table S1a.

Retrospective CT images of the femur heads and jawbones of the MPN patients were provided by the Department of Diagnostic and Interventional Radiology of the RWTH Aachen University with the ethics approval number EK028/19. More detailed information on patient characteristics can be found in Table S1b.

FFPE bone marrow biopsies and peripheral blood plasma of MPN and Healthy donors were provided by the Laboratory of Stem Cells and Cancer Biology, Division Oncology and Hematology, HOCH Health Ostschweiz with the approval from the ethics committee of Ostschweiz (EKOS 23/022). Informed consent was obtained from all participants and/or their legal guardian/s. All samples were retrospectively collected. All MPN patient biopsies were taken from pelvic bones. Healthy donor samples were obtained from either hip surgery or non-hematological malignant samples.

Human iPSCs generated from the peripheral blood mononuclear cells of either healthy or MPN patients were used. iPSC, PDL fibroblast and primary cell work was approved by the ethics committee of the Medical Faculty of RWTH Aachen University (EK206/09, EK374/19).

### Clinical imaging

#### Positron emission tomography/computed tomography (PET/CT)

Radiographic measurements were analyzed retrospectively in MPN patients (N=7) and HD (N=8). A detailed patient cohort is described in the Table S1C. ^18^F-FDG PET/CT scans of MPN patients were acquired as part of a clinical workup to assess for presence of extramedullary disease before alloHSCT at the University Medical Center Hamburg-Eppendorf between 2012 and 2017. Imaging was performed using a standard whole-body protocol with Gemini GXL10 scanner (Philips) as described before^9^. All control subjects were imaged in the Department of Nuclear Medicine at the RWTH Aachen Medical Faculty with a clinical PET/CT system (*i.e.*,

Biograph Vision 600, Siemens Healthineers, Knoxville USA) for suspicion or first staging of lung carcinoma. Under euglycemic conditions 2,5 MBq/kg of FDG were injected in a 0,9% saline solution *via* a peripheral venous catheter. The PET/CT scan was initiated ca. 60 minutes after tracer injection starting with the CT. CT acquisition settings: standard low dose protocol with 35 mA and 120 kVp, with CARE Dose4D and CAREkV enabled to decrease the radiation dose. CT images were reconstructed using a sinogram affirmed iterative reconstruction algorithm (*i.e.*, SAFIRE) to a voxel size of 1,52 × 1,52 × 2,5 mm^3^ in a 512 × 512 matrix. Using a vendor software, CT values were converted into Hounsfield units (HU) using the formula:

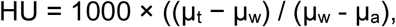

where µ_w_ is the linear attenuation coefficient of water, µ_a_ is the linear attenuation coefficient of air and µ_t_ is the linear attenuation coefficient of the tissue. Immediately after the CT, a static PET acquisition was acquired with an axial field of view of 263 mm using continuous-bed motion (*i.e.*, FlowMotion^®^) with a bed speed of 1,4 mm/s (*i.e.*, equivalent to approximately 100 s/bed position). PET data were reconstructed iteratively using 3-dimensional ordinary Poisson ordered subset expectation maximization (*i.e.*, 3D-OP-OSEM with 4 iterations and 5 subsets) using vendor-recommended settings (*i.e.*, PSF, TRueX and ToF (UltraHD-PET) enabled at zoom of 1x) to a voxel size of 1,65 × 1,65 × 2,50 mm^3^ in a 440 × 440 matrix. Smoothing using a Gaussian filter of 2,00 mm full width at half maximum was automatically applied.

PMOD software package version 4.4 (PMOD Technologies LLC, Zürich, Switzerland) was used to view, render, and analyze the PET/CT data. As organs of interest the following bones were selected and further segmented: the left femoral head, the proximal and distal parts of the left femur, the entire mandible, the entire maxilla, the left humeral head, the proximal and distal parts of the left humerus as well as the proximal part of the tibia. Initially, those regions were manually masked as seen on CT. Afterwards, a CT-based automatic isocontour was generated using +500 HU as the minimal threshold to segment the bone structure. To segment the bone marrow a second automatic isocontour was generated using +500 HU as the maximal threshold. Upon completion of all target volumes of interest, the average, the maximum as well as the average of the top 5 hottest voxels was recorded both for the PET (in standardized uptake values, SUV) and CT data (in Hounsfield units, HU). All VOIs were drawn by a blinded member of our team, in order to exclude analysis bias.

#### Computed tomography (CT)

For the radiological analysis, CT datasets were retrospectively retrieved from the institutional picture archiving and communication system (PACS). A patient cohort with confirmed myeloproliferative neoplasms (MPN; *n* = 34) was analyzed. All available CT examinations were considered, including both non-contrast and contrast-enhanced scans. Image evaluation was consistently performed using bone window settings. Bone density assessment was performed using the PACS software (Philips VUE PACS). Regions of interest were manually placed in the trabecular bone of the femoral neck and the alveolar bone of the mandible and/or maxilla, following a standardized protocol analogous to previously published approaches^12^. Mean attenuation values expressed in Hounsfield units (HU) were recorded for each ROI. Where applicable, multiple measurements per site were obtained and averaged.

#### United Kingdom Biobank (UKBB) study

We analyzed data from the UKBB, a population-based cohort of ∼500,000 participants aged 40–69 years recruited between 2006 and 2010. All participants provided informed consent, and the study was approved by the Northwest Multi-centre Research Ethics Committee.

Analyses were performed under UKB application 245437. MPN was defined using hospital episode statistics from ICD-10 code D47, capturing lifetime-risk for the phenome-wide association analysis (PheWAS), and secondly, we restricted outcomes to musculoskeletal and digestive diagnoses.

#### Murine studies

All animal procedures using JAK2 knock-in model were approved by the Animal Care and Use Committee (German Regierungspräsidium Karlsruhe für Tierschutz und Arzneimittelüberwachung, Karlsruhe, Germany) under approval number G247/18 and G150/23. Mice were maintained in individually ventilated cages under specific pathogen-free (SPF) conditions at the German Cancer Research Center (DKFZ, Heidelberg).

To generate the tamoxifen-inducible JAK2V617FxRosaCreERT1 knock-in mouse model, JAK2-V617F^fl/wt^ mice were crossed with RosaCreERT+/− mice. Littermate pups without the mutation (JAK2WT) were used as control mice. For transplantation experiments, regular C57BL/6J mice (purchased from Janvier Laboratories) were used as recipients. CD45.1+ C57BL/6J background mice were used to create transgenic lines, while competitor C57BL/6J CD45.1+/ CD45.2+ mice were generated by crossing CD45.1+ C57BL/6J mice with C57BL/6J mice.

Thrombopoietin (ThPO) overexpression model in TdTomato+ recipients was used from the previously published cohort^15^.

#### Bone Marrow Transplantations

Recipient mice were lethally irradiated with two rounds of 500 Rad. 24 hours later, mice were transplanted via i.v. injection with 3×10^6^ whole BM cells derived from JAK2V617F;RosaCre or CrexRosaCre donor mice, together with 3×10^6^ WT CD45.1+/CD45.2+ competitive whole BM cells.

#### Flow cytometry of murine hematopoietic cells

Peripheral blood was collected from the facial vein into EDTA-coated tubes and analyzed using a ScilVet abc Plus veterinary blood cell counter (Scil GmbH). Bone marrow was harvested in RPMI + 2% FCS by isolating, cleaning and crushing the bones. Cell suspensions were filtered through a 40 μm cell strainer, centrifuged and resuspended in ACK buffer (Lonza, #10-548E) and Pharm Lyse Buffer for RBC lysis (BD, #555899) for 10 minutes at room temperature (RT). After washing, cell suspensions were stained using following antibodies: CD115-PE-Cy7 (clone AFS98), CD41-APC (clone MWReg30), Ly6C-AF700 (clone HK1.4), CD45.1-APC-Cy7 (clone A20), CD47-BV421 (clone miap301), CD34-PE (clone SA376A4), CD150-PeCy7 (clone TC15-12F12.2), CD117-APC (clone 2B8), CD45.2-AF700 (clone 104), CD3-PB (clone 145-2C11), CD11b (clone M1/70), B220-PB (clone RA3-6B2), Gr1-PB (clone RB6-8C5), Ter119-PB (clone TER119), CD11b-BV711 (clone M1/70) and MHC class II-BV785 (clone M5/114.15.2) from Biolegend; Ly6A/E-PerCP-Cy5.5 (clone E13-161.7), CD48 (clone HM48-1), CD45.1-BV605 (clone A20), CD16/32-BV785 (clone Ab93), CD45.2-PE (clone 104), Ly6G-PE-CF594 (clone 1A8), CD3-BV605 (clone 145-2C11) from BD Biosciences; CD19-PE-Cy5.5 (clone 6D5) from SouthernBiotech. Cells were analyzed by flow cytometry using FACSymphony™ A5 SE Cell Analyzer and LSR Fortessa. Data was analyzed using FlowJo software (FlowJo, LLC, Ashland, OR 97520).

#### *in vivo* treatments

In order to knock in JAK2V617F, primary donor JAK2V617F-LRC mutant mice (JAK2-V617Ffl/wt x RosaCreERT+/− x SCLtTA+/− x H2BGFP+/−) or wild-type Cre-Control mice (JAK2-V617Fwt/wt x RosaCreERT+/− x SCLtTA+/− x H2BGFP+/−) were injected i.p. with 2mg tamoxifen (Merck, #T5648) in 200μL solvent (90% sunflower oil + 10% ethanol). 4 weeks after the last tamoxifen injection, the donor mice were sacrificed by cervical dislocation to collect the cells needed to perform transplantation experiments. For pharmacological treatments, fedratinib (120 mg/kg, HY-10409A, MCE) was dissolved in vehicle solution (0.5% methylcellulose + 0.05% Tween-80 in water) and administered to mice via oral gavage daily, 5 days/week. LSKL (30mg/kg, #HY-P0299, MCE) was dissolved in vehicle solution (0,9% NaCL) and administered via i.p., 3 times/week, for either 4 or 8 consecutive weeks.

#### Histological and immunohistological stainings

Murine organs were fixed in 2% paraformaldehyde for 8h and transferred to 70% ethanol. Murine bones were decalcified in 0.5M EDTA for 7 (femur/tibia) and 21 (jaws) days, dehydrated, and paraffin embedded. H&E was performed according to established routine protocols and reticulin staining was performed using the Reticulin silver plating kit according to Gordon & Sweets (Merck, #1.00251) on 5μm sections.

TRAP staining was performed on FFPE bones using a leukocyte acid phosphatase (TRAP) kit (387A, Sigma-Aldrich, Steinheim, Germany) as per manufacturer’s instructions. The sections were then covered with Aquatex (1.08562.0050, Merck, Darmstadt, Germany).

For immunofluorescence stainings, samples were incubated in blocking solution (10% fetal bovine serum in PBS with 0.1% Triton X-100) for 1 h at room temperature. Primary antibodies with rabbit anti-COMP (Abcam, #ab231977, 1:50 dilution) were diluted in PBS with 1% fetal bovine serum and incubated overnight at 4 °C. Next, slides were washed 3–5 times in PBS in 5 min intervals. Species-specific Alexa Fluor-conjugated secondary antibodies AF-594 (Thermo Fisher Scientific, A11012) diluted 1:500 in PBS with 1% fetal bovine serum and 0.1% Triton X-100 were added and incubated for 1h at RT. Slides were washed 3–5 times in PBS in 5 min intervals. Nuclei were counterstained with DAPI (Sigma-Aldrich, D9542, 1:1,000 dilution). Coverslips were mounted with Entellan (Merck, #1.07961.0100).

For immunohistochemical analysis of Thrombospondin 1 (THBS1) on murine and human (MPN n=7, healthy n=3) FFPE tissues, antigen retrieval was performed using citrate-buffer in a conventional lab microwave (Vector, antigen unmasking solution). Sections were treated with 3% H2O2 and blocked with Avidin/Biotin blocking kit (Vector), and subsequently incubated with primary antibody (polyclonal goat anti-Thbs1, #PA5-142448, Invitrogen, 1:50 for murine and 1:100 for human samples) for 4◦C overnight. Sections were stained with secondary antibodies for 30 min at room temperature. Slides were incubated with AB complex for 30 min at room temperature, washed, and incubated for a further 10 min with DAB substrate. Slides were counterstained with hematoxylin and mounted with glass coverslip. Slides were scanned using a PhenoCycler Fusion system (Akoya Biosciences). Images were exported as qptiff and analyzed using the QuPath software v. 0.5.0.

#### ELISA

ELISAs were performed on human plasma samples (MPN n=293, healthy donor n=74) with human CD47 (DY4670, Biotechne) and TSP1 (Thrombospondin-1, DY3074, Biotechne) DuoSet ELISA Kits as per the manufacturer’s recommendations. Briefly, capture antibody was coated overnight, blocked with blocking buffer (reagent diluent buffer with 1% BSA and 2% FCS) for one hour, 4 times diluted samples were incubated for 2 hours, followed by the detection antibody for 2 hours and finally treated with the Streptavidin-HRP for 30 minutes. After that the TMB substrate was incubated for 20 minutes and 35 minutes for CD47 and TSP1, respectively, the reaction was stopped by adding the stop solution directly to the substrate without any washing step in between. The absorbance at 450nm was measured within 3 minutes after adding the stop solution. CD47 and TSP1 concentrations were calculated from the slopes of the respective standard curves.

#### Murine micro-computed tomography

Femurs, maxillae, and mandibles were fixed in 2% PFA for 8h and scanned in 70% ethanol using a Skyscan 1272 micro-CT system (Bruker Micro-CT, Belgium) at 60 kV and 140 μA with a 0.25 mm aluminum filter, achieving an isotropic voxel size of 5 μm. Data were reconstructed in NRecon software (Bruker Micro-CT, Belgium). For analyses of comparable areas, the data were co-registered to one reference sample in DataViewer (Bruker Micro-CT, Belgium). Subsequently, multiple volumes of interest (VOIs) were defined for quantitative analysis in CTan(Bruker Micro-CT, Belgium).

For femoral specimens, distinct VOIs were established: (1) distal and proximal areas (1.5 mm and 6.5 mm length, respectively), (2) mid-diaphyseal cortical region (0.5 mm length), and (3) growth plate region in the distal femoral head (cylindrical VOI, 0.4 mm diameter × 1.0 mm height). The trabecular part of the femur was defined virtually using a cascade of opening and closing algorithms. The cortical compartment was defined through ROI shrink-wrap algorithms following trabecular bone subtraction. Bone mineral density (BMD) values were calibrated using hydroxyapatite phantoms scanned under identical conditions. Cortical thickness was estimated after the subtraction of the trabecular region and using an identical algorithm for definition of trabecular thickness. Growth plate thickness was quantified within the cylindrical VOI using the same morphological approach. Femoral length measurements were obtained from the co-registered datasets.

Analysis of maxillary and mandibular specimens were performed in several VOIs: (1) alveolar bone of the first molar tooth socket (M1; 0.8 mm height in maxilla, 0.6 mm height in mandible), (2) alveolar bone between the first and the second molars (M1-2), (3) alveolar bone around the third molar (M3), and (4) periodontal ligament space of the first molar (PDL-1). Bone and periodontal ligament parameters were quantified using established algorithms as previously described^37^. For bone loss assessment, total bone volume was analyzed, incorporating both vertical bone loss and increased alveolar porosity. Periodontal ligament space thickness was determined using standardized trabecular thickness algorithms.

### Sequencing

#### FACS sorting and library preparation

Murine jaws were crushed and flushed using PBS/2%FCS and filtered through a 70µm nylon mesh. The remaining bone chips were digested in 1mg/mL Collagenase II (Thermo Scientific, #17101-015) and DNAse I (100µg/mL, Sigma-Aldrich) at 37 ◦C for 30 minutes. After pooling of the supernatant of the bone chips with the flushed fraction, cells were were stained with the following antibodies: CD45-APCCy7 (clone 30-F11, BD), Ly6A/E-PerCP-Cy5.5 (clone E13-161.7, Biolegend), TER119-PB (clone TER119, Biolegend) and DAPI. Murine jawbone cells were sorted based on the following expression patterns: DAPI-TER119-CD45+ (hematopoietic cells), DAPI-TER119-CD45-Ly6A/E+ (stroma). Following sorting, the sorted populations were mixed at variable ratios, on a sample-to-sample basis, to ensure recovery of all main cell populations. The resulting cell suspension was centrifuged, resuspended in PBS/2%FCS and processed for scRNAseq (10X Genomics, protocol CG000315) using the manufacturer’s procedure.

Patient iliac crest biopsies were crushed and flushed using PBS/2%FCS and filtered through a 70µm nylon mesh. The remaining bone chips were digested in Liberase TL (125µg/mL, Roche, for iliac crest biopsies) or 3mg/mL Collagenase I (LS004196, Worthington Biochemical, for teeth samples) and DNAse I (100µg/mL, Sigma-Aldrich) at 37 ◦C for 30 minutes, with gentle agitation. Following this incubation, supernatant was collected and strained through a 70µm nylon mesh. Bone chips were additionally washed twice with PBS/2%FCS before pooling it with the flushed fraction.

Isolated cells were resuspended in PBS/2%FCS and stained at 4 ◦C for 20 minutes using the following antibodies: CD34-PE (BD, #555822), CD45-APC (BD, #555485), CD3-PerCpCy5.5 (BD, #560835), CD19-BB700 (BD #566396), HLA-DR-PE-Cy7 (BD, #560651), CD235ab-Pacific Blue (BioLegend, #306612), CD45-BV786 (BD, #664535), CD235a-BV421 (BD, #562938). Then, cells were diluted in PBS/2%FCS and stained with Calcein Green (#C34852 Invitrogen, 1µM final) for 10 minutes at room temperature. After washing, cells were resuspended in 400µL of PBS/2%FCS and sorted using a BD FACS Melody. Cells from the iliac crest biopsies were sorted based on the following expression patterns: CD34+ (HSC-enriched), CD34− CD45− CD235a− (stroma-enriched) and CD34− CD45+ HLA-DR+ CD3-CD19− (mononuclear phagocyte-enriched). Cells from the human teeth samples were sorted based on the following antibodies: DAPI-CD235a-CD45+ (hematopoietic cells), DAPI-CD235a-CD45− (stroma). Following sorting, the sorted populations were mixed at variable ratios, on a sample-to-sample basis, to ensure recovery of all main cell populations. The resulting cell suspension was centrifuged, resuspended in PBS/2%FCS and processed for scRNAseq (10X Genomics, protocol CG000315 and CG000731) using the manufacturer’s procedure.

All libraries were prepared using the Chromium Next GEM Single Cell 3’ Reagent Kits (v3.1 and v4): Library Construction Kit and Gel Bead Kit (PN-1000268), Chromium Next GEM Chip G Single Cell Kit (PN-1000127) and Dual Index Kit TT Set A (PN-1000215), and following the Chromium Next GEM Single Cell 3ʹ v3.1 and v4 (Dual Index) User Guide (CG000315, Rev F and CG000731, Rev B). Finalized libraries were sequenced on a Novaseq6000 platform (Illumina), aiming for a minimum of 20.000 reads/cell using the 10x Genomics recommended number of cycles (28-10-10-90 cycles).

Raw counts were aligned to mouse or human genome (mm10 or hg18) and quantified using 10x Genomics Cell Ranger (version 7.0.0).

#### Single-cell RNA sequencing analysis

Single-cell RNA-sequencing data were processed using R (v.4.4.1) and Seurat (v.5.3.1). Cells failing general quality control (QC) thresholds, including low gene counts, low UMI counts, high mitochondrial gene content (>30%), or >50% ambient RNA (using decontX; Celda v 1.14.2), were excluded from downstream analyses.

After QC filtering, a total of 95,368 cells from human bone marrow and teeth samples were retained for analysis. For the mouse dataset, samples from the maxilla and mandible contained a total of 15,790 cells for downstream analysis. Data integration across samples was performed using Harmony (v.1.2.4) to correct for batch effects and doublets were identified and removed using DoubletFinder (v. 2.0.6). The integrated dataset was visualized using UMAP. Cell type annotation was performed manually based on the expression of known marker genes. Subclustering was conducted for selected populations when higher resolution was required. Cell type prioritization between conditions was performed using Augur (v. 1.0.3). Differential gene expression analysis between control and diseased samples was carried out in a pseudobulk aggregation approach with DESeq2 (v.1.44) using the Wald test. Genes identified as differentially expressed were subjected to pathway enrichment analysis using Reactome (R.), Gene Ontology (GO), WikiPathways (WP), KEGG, BioCarta, and PID databases. Enrichment significance was evaluated using the fgsea permutation-based enrichment test. Differences in gene and module scores expression between conditions were tested using an unpaired two-sided Wilcoxon rank-sum test, and significance was reported using p-values (non-significant comparisons not shown).

Cell–cell communication analyses were performed using CellChat, and interactions were filtered to those supported by ≥10 cells per group. Statistical significance was assessed using bootstrap resampling (nboot = 100). Global cell–cell interaction changes were summarized using heatmaps based on the delta probability of interaction. Selected cell–cell interactions were visualized using alluvial plots, in which the delta interaction probability was explicitly represented to highlight condition-specific signaling changes. For the human dataset, an intra-patient comparison was conducted to identify differences in signaling between bone marrow and teeth samples. For these analyses, delta–delta interaction probabilities were calculated and reported to quantify tissue-specific changes in signaling.

For the mouse dataset, trajectory inference was performed using Slingshot (v.2.12.0), with inferred lineages visualized by overlaying trajectories on LAP reduction plots. Condition-specific dynamics were further visualized using stream plots, with pseudotime values normalized across conditions. Transcription factor activity analysis was performed using decoupleR (v.2.10.0), leveraging the CollecTRI database to infer regulatory activity. Gene module scores were computed for predefined gene sets curated from the literature.

#### Spatial transcriptomics

Visium CytAssist spatial transcriptomics data from tibial sections were obtained from our previously published work^15^ and reanalyzed in this study. Experimental procedures, including FFPE section processing, hematoxylin–eosin staining, image acquisition, library preparation, and sequencing on an Illumina NovaSeq platform, were performed as previously described. Brightfield images were aligned to capture areas, and segmentation files were generated using Loupe Browser (v8.1.2, 10x Genomics). Raw sequencing data were processed with Space Ranger v3.0.0 to generate spot-level gene expression matrices.

Downstream analysis was performed in Python (v3.10.16) using Scanpy (v1.11.0) and Squidpy (v1.6.5). Spots with 200–7,500 total counts, mitochondrial transcript content >5%, or genes detected in fewer than 10 spots were excluded. Spatial neighbors were computed for each replicate, and Moran’s I was used to identify genes with significant spatial autocorrelation (normalized p < 1×10⁻⁴) shared across all replicates. The filtered data were normalized to 10,000 counts per spot, log-transformed, scaled, and subjected to PCA. A k-nearest neighbor graph was constructed using 20 principal components and 15 neighbors, followed by Leiden clustering (resolution = 0.3). Differentially expressed genes were identified using a two-sided Wilcoxon rank-sum test, and clusters were mapped back onto tissue sections and annotated based on highly expressed genes.

### In vitro studies

#### Immortalization and characterization of PDL cell lines

Human PDL cells (mandible, molar 38) were isolated from a 32 y.o. female patient. Primary human periodontal ligament (hPDL) cells were cultured from periodontal connective tissue, isolated from the middle root section of healthy human teeth. Only decay-free teeth from the patient, which needed to be extracted for medical reasons, were used for human PDL cell isolation. Immediately after extraction, the teeth were transferred into an isolation medium of DMEM high-glucose (Gibco, Billings, MT, USA), 10% FetalBovine Serum (FBS), qualified heat-inactivated (Gibco), 50 mg/L ascorbic acid (Sigma, Saint Louis, MO, USA), antibiotic-antimycotic (Gibco) and stored at 4◦C until isolation was started within 24 h after extraction. For isolation, The PDL tissue, which is located on the lower three-fifths of the height of the root, was then scraped off and incubated in a solution of 2 mg/mL collagenase type I (Worthington Biochem, Freehold, NJ, USA) for 1.5 h at 37◦C. After incubation, cells were plated into 6-well plates. After the first splitting, the isolation medium was replaced by a culture medium of DMEM high glucose (Gibco), 10 % FBS (Gibco), 50 mg/L ascorbic acid (Sigma), Penicillin/Streptomycin (Gibco). Primary PDL cells were immortalized with hTERT and SV40 Large T at passage 2. For osteogenic differentiation, cells were seeded in 24-well culture plates in a density of 10.000/cm^2^ for osteogenic differentiation. The osteogenic induction medium consists of DMEM low glucose (Fisher Scientific, Schwerte, Germany), 10% FCS, 100 nM dexamethasone, 10 mM sodium-glycerophosphate, and 0.05 mM L-ascorbic acid (all from Sigma-Aldrich, Taufkirchen, Germany). Three times per week, a medium exchange was performed. For the microfluidic experiment, 20.000 of immortalized PDL fibroblasts were seeded in each well of a 96-well plate and centrifuged using 350g 5 min to form spheroids before chip loading.

#### Maintenance of induced pluripotent stem cells

Human induced pluripotent stem cells (iPS cells) were maintained on growth factor–reduced basement membrane matrix (Geltrex™, Gibco, #A1569601) in StemFlex™ basal medium (Gibco, #15627578) supplemented with 1x penicillin/streptomycin (PAN-Biotech, #P06-07100) at 37 °C and 5% CO₂. Medium was exchanged daily, and cells were passaged at 60–80% confluency either as single cells using Accutase (Pan Biotech, #P10-21500) or as small aggregates using EDTA-based dissociation. For single-cell passaging, the ROCK inhibitor Y-27632 (10 µM; Abcam, #129830-38-2) was added for the first 24 h after seeding to enhance cell survival.

#### Viral transduction of GFP and RFP for iPS cells

MPN patient–specific induced pluripotent stem (iPS) cell lines (JAK2^V617F/V617F^ and CRISPR repaired isogenic JAK2^WT/WT^, patient 2 with PV diagnosis, JAK2V617F variant allele frequency 96%) were generated as previously described^38,23^. GFP or RFP labeling of iPS cell lines was achieved using GFP- or RFP-expressing pWPXL lentiviral vectors produced in HEK293T cells with the psPAX and pMD2.G packaging plasmids. iPS cell cultures at 60–70% confluency were incubated overnight at 37 °C in a humidified atmosphere containing 5% CO₂ with a 1:1 mixture of iPS-Brew medium and viral supernatant supplemented with 8 µg/mL polybrene. Following transduction, iPS cells were expanded and sorted for GFP⁺ or RFP⁺populations using a BD FACSMelody™ cell sorter. Sorted cells were cultured on Laminin-521–coated plates (BioLamina) in the presence of a Rho kinase inhibitor (MedChem Express) for 24 h, followed by further expansion on Matrigel-coated plates.

#### Differentiation of iPSC-derived CD34+ cells

MPN patient–specific iPSC-derived hematopoietic progenitors were generated using a modified spin embryoid body (SpinEB) protocol as previously described^23,38,39^. Briefly, single-cell–dissociated iPS cells were aggregated by centrifugation in serum-free differentiation medium composed of IMDM and Ham’s F12 (1:1; Gibco, #12440053 and #11765054) supplemented with 0.4% bovine serum albumin (home-made), chemically defined lipids (Gibco, #11905031), 2 mM GlutaMAX (Gibco, #35050061), 0.4 mM monothioglycerol (Sigma-Aldrich, #96-27-5), 50 µg/mL L-ascorbic acid (Stemcell Technologies, #72132), and 6 µg/mL human transferrin (Sigma-Aldrich, #11096-37-0). Mesoderm and hematopoietic specification was induced by supplementation with 10 ng/mL FGF2 (PeproTech, #100-18B), 10 ng/mL BMP4 (Miltenyi Biotec, #130-111-168), and 10µM Y-27632 ROCK inhibitor (Abcam, #129830-38-2). From day 2 until 14, EBs were cultured with daily partial medium exchange and stage-specific cytokine supplementation, including 1× stem cell factor (Miltenyi Biotec, #130-096-692), 25 ng/mL VEGF (Miltenyi Biotec, #130-109-385), 10 ng/mL FLT3L (PeproTech, #300-19-250), 30 ng/mL IL-3 (ImmunoTools, #11340037), 25 ng/mL hyper–IL-6 (Stem Cell Technologies, #78050) and 50ng/mL M-CSF (Thermo Fischer, #300-25).

#### Bone marrow organoids culture

BM organoids were generated using the newly established HD353.6 line based on previously described protocol^22^. In brief, for mesoderm induction, cell aggregates were cultured in phase I medium composed of STEMDiff APEL2 medium (StemCell Technologies, #05275) supplemented with 50ng/mL FGF2 (PeproTech, #100-18B) and 50ng/mL BMP4 (Miltenyi Biotec, #130-111-168), 3mM CHIR99021 (Merck, #SML1046) and 25ng/mL VEGF (Miltenyi Biotec, #130-109-385). For hematopoietic and vascular commitment, organoids were cultured in phase II medium consisting of 10ng/mL SCF (Miltenyi Biotec, #130-096-692), 25ng/mL FLT3L (Peprotech, #300-19-250), 25ng/mL VEGF-A (Miltenyi Biotec, #130-109-385). From day 5 onward, organoids were embedded in a collagen-based hydrogel containing 60% collagen (collagen type I (Corning, #354236), collagen type IV (Cell Systems, #5022)) and 40% matrigel (Corning, #354230) and cultured under vascular sprouting conditions using phase III media supplemented with additional 5% KO serum replacement (Gibco, #10828010), 5U/mL heparin (Merck, #H3149), 50ng/mL FLT3L (PeproTech, #300-19-250), 20ng/mL IL-3 (ImmunoTools, #11340037), 20ng/mL hyper–IL-6 (Stem Cell Technologies, #78050), 50ng/mL M-CSF (PeproTech, #300-25), 50ng/mL TPO (Peprotech, #315-14) and 50ng/mL EPO (Immunotools, #11344795). From day 13 onward, organoids were maintained in phase IV maturation medium containing the same cytokine composition, enabling the development of functional stromal, vascular, and hematopoietic compartments. Media were replenished every 72 hours. For drug treatment, samples were treated with vehicle (water) or LSKL (5µM, #HY-P0299, MCE) either statically or added on top of the fluidic unit starting from 7 days post engraftment of exogenous cells for three times with media replenishment every second day.

#### Engraftment of exogenous cells into bone marrow organoids

On day 14 of SpinEB protocol (see above), EB-derived cells from JAK2V617F;RFP and JAK2WT;GFP iPSC lines were harvested and CD34⁺ cells were enriched by magnetic-activated cell sorting using CD34 UltraPure MicroBeads (Miltenyi Biotec, #130-100-453) for subsequent use as donor cells for bone marrow organoids. Primary patient-derived CD34+ HSPCs were isolated from the peripheral blood mononuclear cells using CD34 UltraPure MicroBeads (Miltenyi Biotec, #130-100-453) and stained with CellVue (Sigma Aldrich, #12352207) as per manufacturer’s instructions.

#### Organ-on-a-chip

Microfluidic experiments were performed using an Ibidi pump system (Ibidi, #10905) and fluidic units (Ibidi, #10903) with µ-slide III 3D perfusion chips (Ibidi, #80376), perfusion set black (Ibidi, #10966), serial connectors (Ibidi, #10830). Bone marrow organoids containing exogenous CD34+ iHSPCs were loaded into the central gel channel of microfluidic chips and co-cultured with PDL fibroblast spheroids placed in adjacent wells. Chips were connected to a perfusion system and cultured under continuous unidirectional flow using organoid culture medium at 37 °C and 5% CO₂ for 7 days.

#### qRT-PCR

Total RNA from whole BM organoids or PDL spheroids was extracted using PureLink RNA Mini kit (Thermo Fischer, #12183018A) and reverse-transcribed using the iScript™ cDNA Synthesis Kit (Bio-Rad; #1708890) as per kit instructions. qPCR was performed on a CFX Opus 96 Real-Time PCR system (Bio-Rad) using a hard-shell 96-well PCR plate (Bio-Rad; #HSP9601). Gene expression was analyzed using primers for *ACTA2* (forward 5′-TCCTTCATCGGGATGGAGTCT-3′, reverse 5′-TACATAGTGGTGCCCCCTGA-3′), *GAPDH* (forward 5′-GAAGGTGAAGGTCGGAGTC-3′, reverse 5′-GAAGATGGTGATGGGATTTC-3′), *IL1β* (forward 5′-CACTTACATTGTCACCAGAGAACC-3′, reverse 5′-ATCGTGCACATAAGCCTCGT-3′), *THBS1* (forward 5′-GACGCCTGCCCCATCAAT-3′, reverse 5′-TCTGTACCCCTCCTCCACAG-3′), *CYTL1* (forward 5′-TCGGAGCCCTCGTTGGA-3′, reverse 5′-TCCTCCACAATGCTACGGC-3′), *COL1A1* (forward 5′-ACGGCTGCACGAGTCACAC-3′, reverse 5′-GGCAGGCGGGAGGTCTT-3′), *CD47* (forward 5’-CCTGAAATCAGAAGAGGGCCA-3’, reverse 5’-AGTCTCTGTATTGCGGCGTG-3’), *ACAN* (forward 5’-TGGGAACCAGCCTATACCCCAG-3’, reverse 5’-TGGGAACCAGCCTATACCCCAG-3’) and *SOX9* (forward 5’-TCTCGCTTCAGGTCAGCCTTG-3’, reverse 5’-GCATGAGCGAGGTGCACTC-3’). Relative gene expression was calculated using the ΔΔCt method with *GAPDH* as the reference gene.

#### Flow cytometry of human *in vitro* experiments

For assessment of the hematopoietic compartment in collagenase IV–digested BMOs (Collagenase IV, Gibco, #17104019), antibodies against CD45 (BV786, clone HI30, BD Biosciences, #563716), CD14 (BV605, clone 63D3, BioLegend, #367126), CD235a (Pacific Blue, clone HIR2, BioLegend, #306612), CD16 (BB700, clone B73.1, BD Biosciences, #742286), CD61 (PE-Cy7, clone VI-PL2, BioLegend, #336415), CD34 (PE, clone 581, BD Biosciences, #555822), CD31 (APC-Cy7, clone WM59, BioLegend, #303120), and CD66b (APC, clone G10F5, BioLegend, #312115) were used. Donor origin was determined by endogenous GFP or RFP fluorescence of fluorescently tagged iPSC line.

#### Staining of *in vitro* cultures

Alizarin Red staining was used to visualize calcium-rich deposits developed during osteogenic differentiation of PDL fibroblasts. The cells were fixed in ice-cold 70% ethanol (−20 °C) for one hour and stained with 40 mM Alizarin Red S (Sigma-Aldrich, Taufkirchen, Germany) for 10 min. PBS was used to wash away unbound Alizarin Red, and microscopic images were recorded.

For immunofluorescence analysis of iPSCs, cells on coverslips were fixed with 4% paraformaldehyde, permeabilized with 0.3% Triton X in PBS and blocked with normal goat serum. Cells were incubated with primary antibodies against SSEA4 (mouse monoclonal, Invitrogen, #41-4000, 1:75), NANOG (rabbit polyclonal, Abcam, #ab21624, 1:75), TRA-1-81 (mouse monoclonal, Abcam, #ab16289, 1:250), OCT4 (rabbit polyclonal, Abcam, #ab19857, 1:300), SOX17 (goat polyclonal, R&D Systems, #AF1924-SP, 1:150), NCAM-1/CD56 (goat polyclonal, R&D Systems, #AF2408, 1:150), and β-tubulin III (mouse monoclonal, Sigma-Aldrich, #T8660, 1:1000) overnight at 4 °C. Primary antibody binding was detected using species-appropriate secondary antibodies with 1:500 dilution, including Alexa Fluor 546–conjugated goat anti-mouse IgG (Invitrogen, #A11003), Alexa Fluor 488–conjugated goat anti-rabbit IgG (Invitrogen, #A11008), Alexa Fluor 546–conjugated goat anti-mouse IgG (Invitrogen, #A21045), and Alexa Fluor 488–conjugated donkey anti-goat IgG (Invitrogen, #A11055) and DAPI. Dried coverslips were mounted with Entellan (Merck, #1.07961.0100). Fluorescence was observed and pictures were taken with Leica Stellaris 5 LIA.

### Statistical analysis

Statistical analysis, excluding that for scRNaseq data and UKBB study, was conducted using GraphPad Prism Version 10. Unless otherwise stated in the figure legends, two-tailed unpaired Student’s *t* test was used for statistical analysis. Data are shown as data ± SEM, and a P-value of less than 0.05 was considered statistically significant. *P < 0.05; **P < 0.01; ***P < 0.001; ****P < 0.0001.

