## Supplementary Figures 1-6 for "Oncodevelopmental plasticity of the skeleton in myeloid neoplasms"

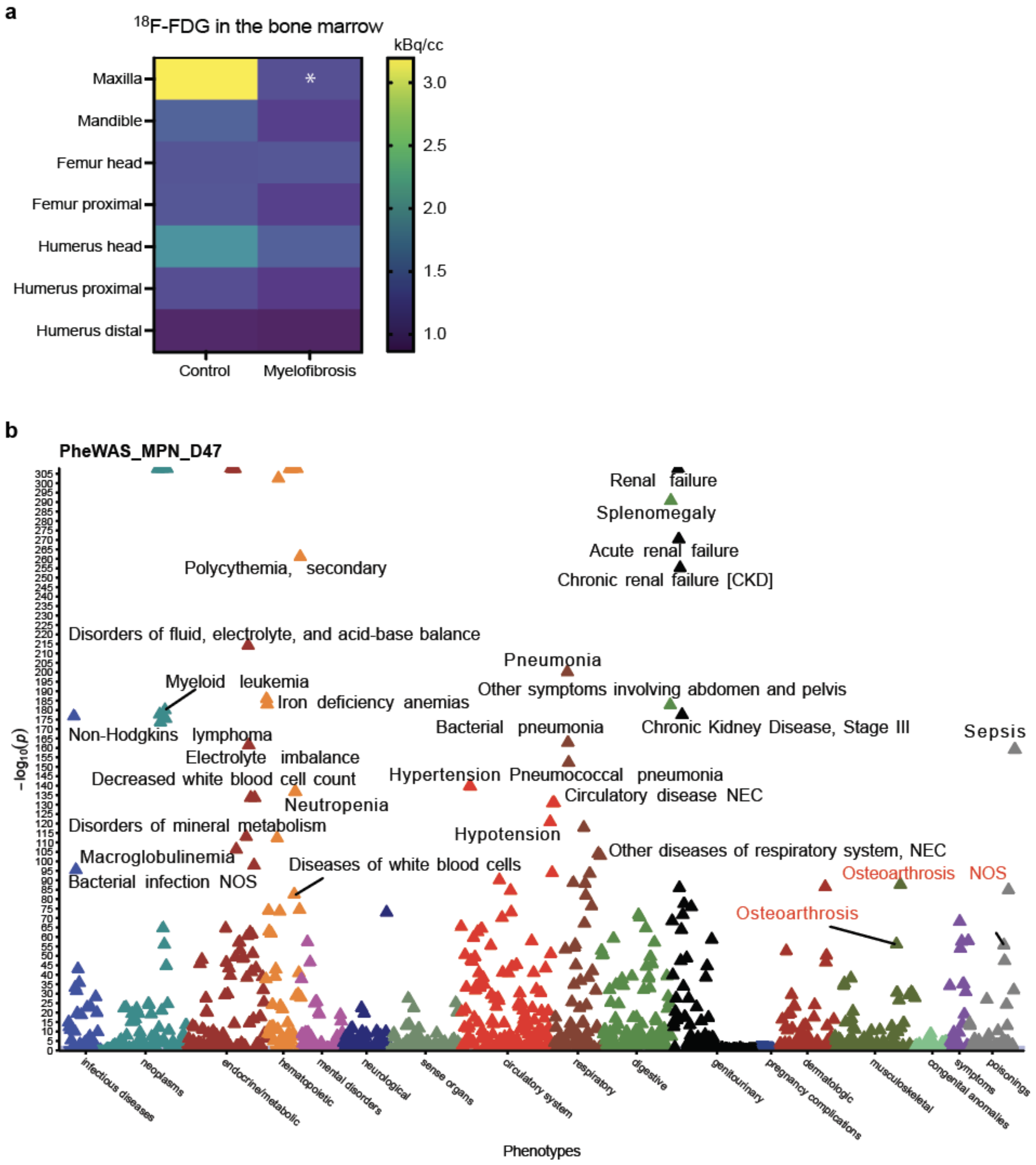


**Fig. S1 (related to Fig. 1): Preferential periodontal bone involvement across ontogenetically distinct skeletal sites in MPN patients.**

**a,** Systemic assessment of the metabolic activity across distinct skeletal sites comparing MF (n=7) versus control (n=8) patients. Two-way ANOVA followed by Sidak's multiple-comparisons test. **b,** Phenome-wide association study from UK Biobank with all phenotypes highly associated with MPN individuals (n=2,548; ICD-10 D47).

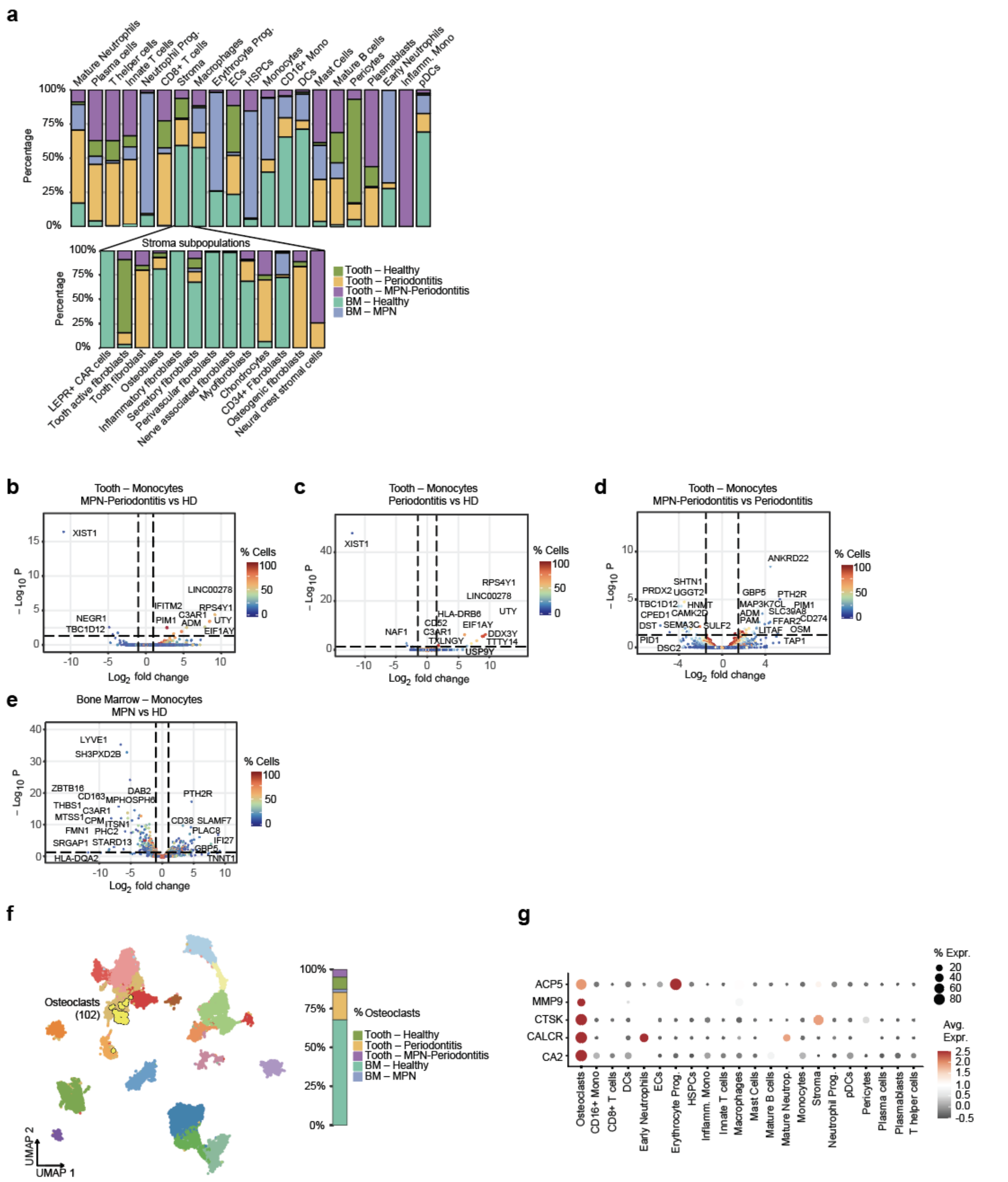

**Fig. S2 (related to Fig. 2): Intrapatient single-cell mapping of ontogenetically distinct skeletal niches uncovers cellular programs in myelofibrosis driving pathologic bone loss.**

**a,** Contribution of annotated cellular clusters per tissue and condition. **b-e,** Volcano plot of differentially expressed genes (DEGs) of monocytes comparing the expression of genes between MPN-Periodontitis versus healthy donor (HD) **(b)**, Periodontitis without MPN versus healthy donor **(c)**, MPN-Periodontitis versus Periodontitis without MPN **(d)** in the teeth as well as MPN versus healthy donor **(e)** in the bone marrow biopsies. **f,** UMAP representation of the integrated single-cell transcriptomic dataset showing annotation of osteoclast subpopulation (n=102 cells) within a macrophage cluster, including the contribution of this subcluster to each condition and tissue. **g,** Dot plot showing top 5 osteoclast-specific markers being highly expressed by the osteoclast cluster.

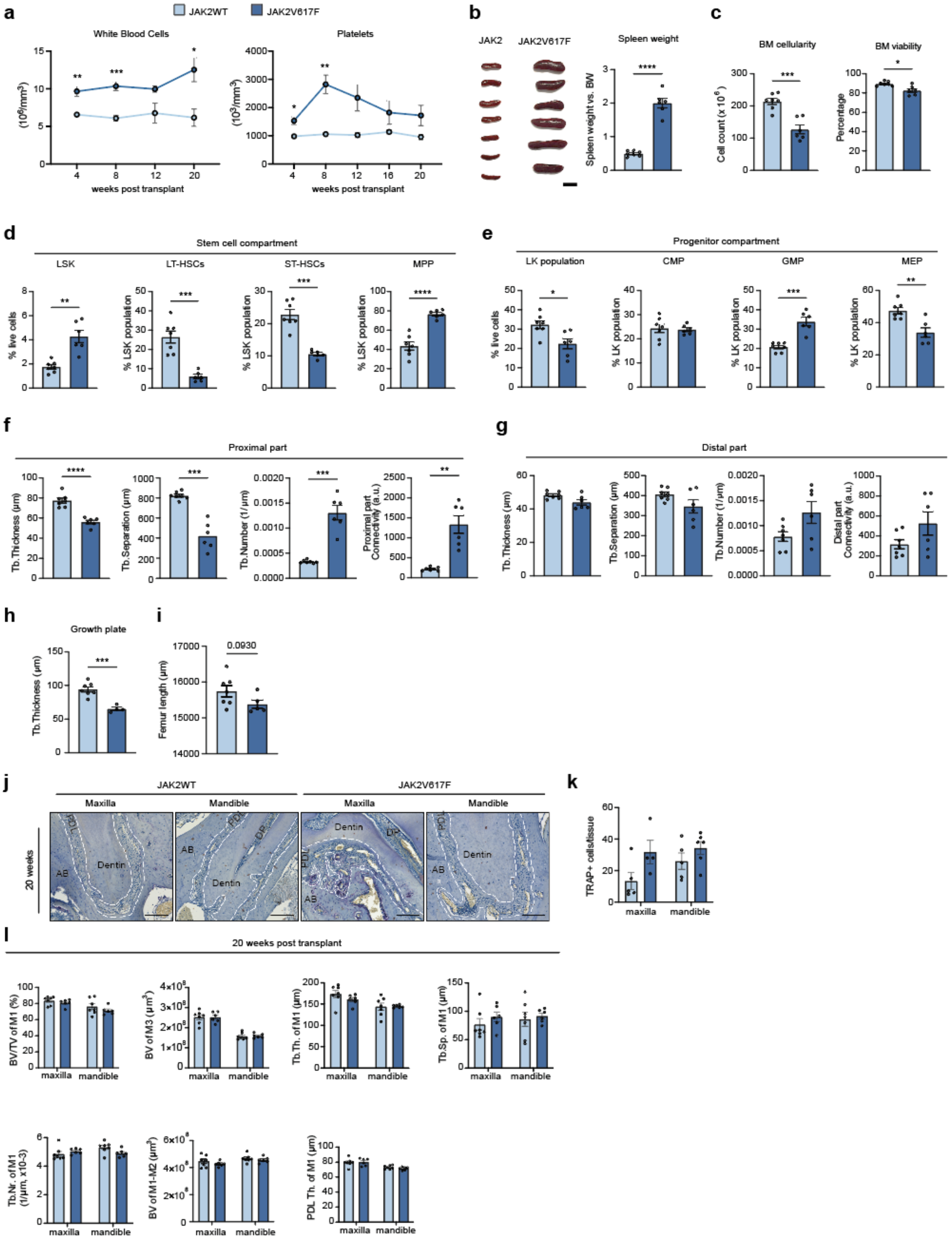


**Fig. S3 (related to Fig. 3): JAK2V617F-driven myelofibrosis drives co-existing osteosclerosis and bone loss in ontogenetically distinct bones.**

**a,** Longitudinal blood counts of experimental mice (n=6 mice per group) used for micro-computed tomography (µCT). Data were analyzed by two-way ANOVA followed by Sidak's multiple-comparisons test. **b,** Spleen size and spleen weight versus body weight (BW) at sacrifice. Scale bar, 1cm. **c,** Cell counts of the bone marrow (BM cellularity) and frequency of viable cells (BM viability) at the end of the experiment. **d,** Flow cytometric quantification of the hematopoietic stem cell (HSC) compartment from the mesoderm-derived bones at the time of sacrifice, including Lin-Sca1+cKIT+ (LSK), long term HSCs (LT-HSCs), short term HSCs (ST-HSCs) and multipotent progenitors (MPPs), in JAK2WT and JAK2V617F mice (n = 6). **e,** Flow cytometric quantification of the hematopoietic stem and progenitor cell (HSPC) compartment from the mesoderm-derived bones at the time of sacrifice, including Lin-Sca1-cKIT+ (LK), common myeloid progenitor (CMP), granulocyte-macrophage progenitor (GMP), megakaryocyte-erythrocyte progenitor (MEP) in JAK2WT and JAK2V617F mice (n = 6). **f-h,** Quantification of the trabecular (Tb.) parameters in the proximal part **(f)**, distal part **(g)** and growth plate **(h)** of the femur from JAK2WT and JAK2V617F mice (n = 6 per group) using 3D reconstructed µCT images. **i,** Quantification of the femoral length using 3D reconstructed µCT images. **j,** Representative images of the TRAP staining on the murine jaws at 20 weeks post transplant. PDL: periodontal ligament, DP: dental pulp, AB: alveolar bone. Scale bar, 50µm. **k,** Quantification of TRAP+ cells per tissue area. Two-way ANOVA followed by Sidak's multiple-comparisons test was used. **l,** Quantification of bone volume (BV) and trabecular thickness (Tb.Th), separation (Tb.Sp.), number (Tb.Nr.) as well as periodontal ligament thickness (PDL Th.) in the jawbones encompassing molar 1 (M1) and the inter-molar region between molars 1 and 2 (M1–M2).

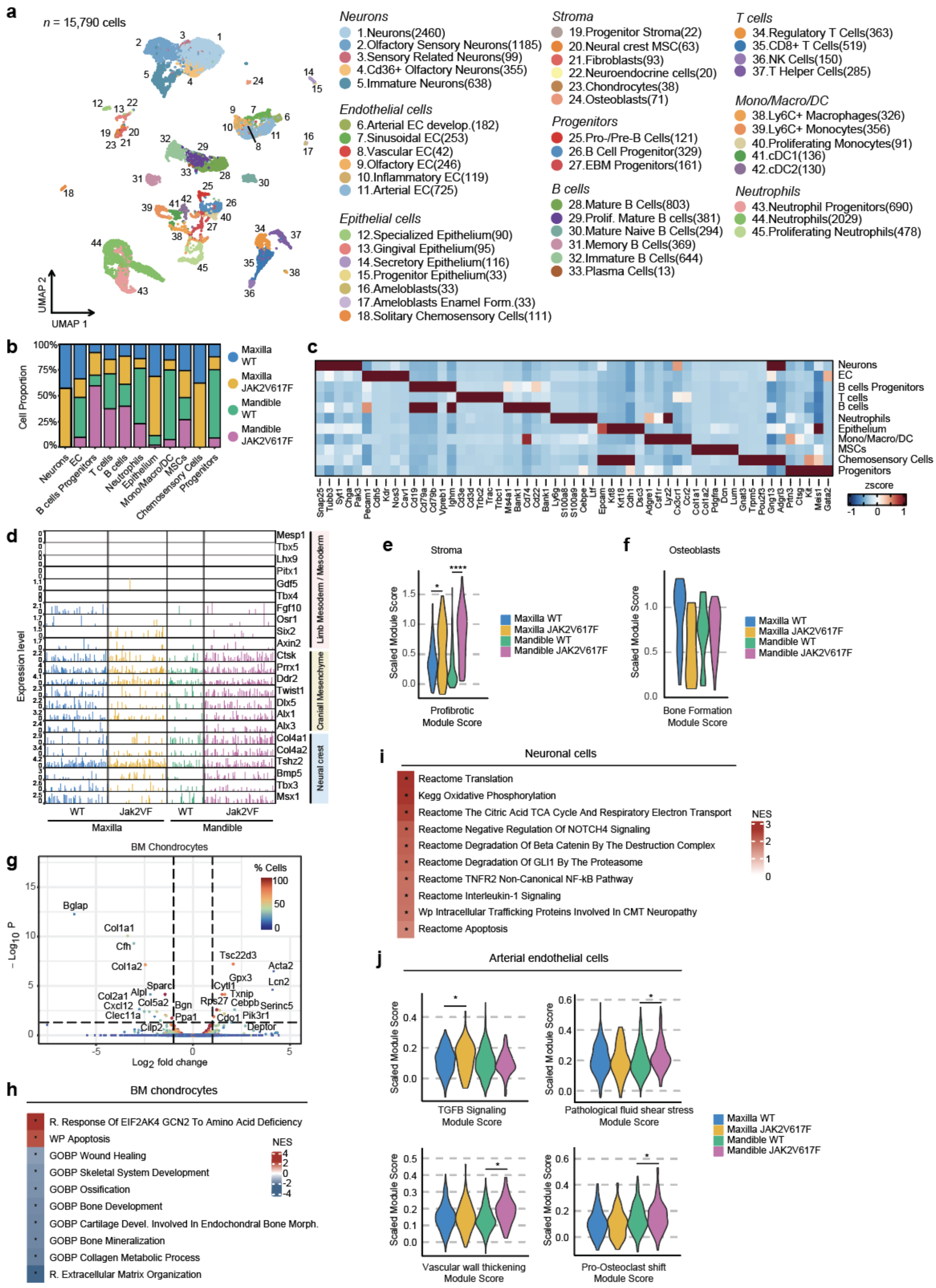


**Fig. S4 (related to Fig. 4): MPN causes the lineage plasticity of neural crest-derived stromal cells driving ectopic chondrogenic fate and contributing to bone loss.**

**a,** UMAP representation of the integrated single-cell transcriptomic dataset from two jawbones in both conditions, comprising 15,790 cells resolved into 45 transcriptionally distinct subclusters (granular annotation). **b,** Contribution of annotated cellular clusters (broad annotation) per condition. **c,** Average gene expression of top 5 marker genes within major cell populations. **d,** Tracksplot of neural crest, cranial mesenchyme and mesoderm marker genes as found in maxilla and mandible in both JAK2WT and JAK2V617F conditions. **e,** Violin plots indicating module score for profibrotic gene set in the neural-crest derived stromal cells per condition. **f,** Violin plots indicating module score for bone formation gene set in the neural-crest derived osteoblasts per condition. **g,** Volcano plot of differentially expressed genes (DEGs) of chondrocyte population in the mesoderm derived bones (femur, tibia, hip bone, spine) comparing the expression of genes between fibrotic (thrombopoietin, ThPO) versus control (empty vector, EV) conditions. **h,** Pathway enrichment analysis for mesoderm-derived chondrocytes comparing fibrotic (ThPO) versus control (EV) conditions. Pathway analysis was performed using Reactome (R.), Gene Ontology (GO), WikiPathways (WP), KEGG, BioCarta, and PID databases. **i,** Pathway enrichment analysis for neurons comparing JAK2V617F versus JAK2WT in maxilla. Pathway analysis was performed using Reactome (R.), Gene Ontology (GO), WikiPathways (WP), KEGG, BioCarta, and PID databases. **j,** Violin plots indicating module scores for indicated signatures in the arterial endothelial cells per condition.

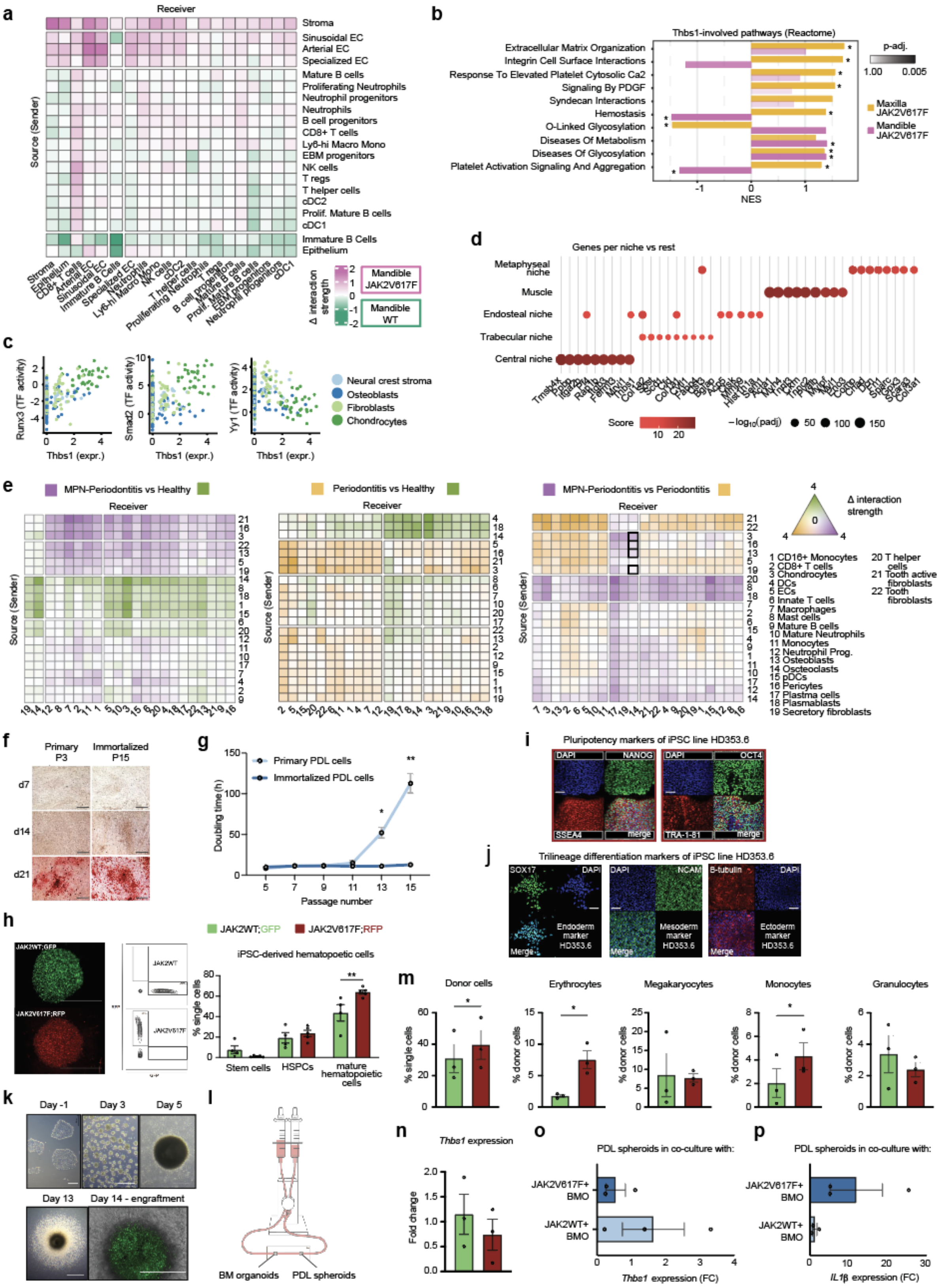


**Fig. S5 (related to Fig. 5): A conserved THBS1⁺ stromal population mediates injury responses across ontogenetically distinct bones in myelofibrosis.**

**a,** CellChat-based intercellular communication networks in paired mandibular jawbones comparing JAK2V617F versus JAK2WT conditions. **b,** Top 10 THBS1-involved pathway enrichment analysis comparing JAK2V617F versus JAK2WT in maxilla and mandible.

Pathway analysis was performed using Reactome (R.). **c,** Transcription factor (TF) analysis on the murine neural crest-derived stromal subclusters showing positive and negative correlation of TF and normalized *THBS1* expression in cells. **d,** Average gene expression of marker genes within 5 annotated spatial domains. **e,** Intercellular communication networks in the patients' dental biopsies comparing color-coded conditions. **f,** Alizarin red staining on primary (passage 3, P3) or immortalized PDL fibroblasts (passage 15, P15) on day 7, 14 and 21 of osteogenic differentiation. Scale bar, 500µm **g,** Doubling time in hours demonstrating proliferation of primary and immortalized PDL fibroblasts upon passages. Two-way ANOVA followed by Sidak's multiple-comparisons test was used. **h,** Immunofluorescence image and representative FACS plot of JAK2V617F;RFP (red) and isogenic JAK2WT;GFP (green) iPS cells and their directed differentiation into hematopoietic lineage. Scale bar, 1000µm. Two-way ANOVA followed by Sidak's multiple-comparisons test was used. **i,** Immunofluorescence staining of pluripotency-associated markers on healthy donor-derived HD353.6 iPSC line. Scale bar, 50µm. **j,** Immunofluorescence staining of trilineage differentiation markers (endoderm, mesoderm, ectoderm) on healthy donor-derived HD353.6 iPSC line. Scale bar, 50µm. **k,** Representative images of HD353.6 iPSC-derived bone marrow organoid differentiation, maturation and engraftment with exogenous fluorescently tagged cells. Scale bars, 500µm and 1000µm. **l,** Schematic overview of the microfluidic organ-on-a-chip model co-culturing bone marrow (BM) organoids with periodontal ligament (PDL) fibroblast spheroids. **m-n,** Flow cytometric analysis of donor-derived hematopoietic cells **(m)** and qRT-PCR of *THBS1* expression **(n)** from BM organoids 7 days after co-culture with PDL fibroblast spheroids. Red bars represent JAK2V617F;RFP-derived cells, green bars represent JAK2WT;GFP-derived cells. **o-p,** qRT-PCR of *THBS1* **(o)** and *IL1β* **(p)** expression in PDL fibroblast spheroids 7 days after co-culture on the microfluidic chip with BM organoids (either engrafted with JAK2V617F;RFP or JAK2WT;GFP iHSPCs).

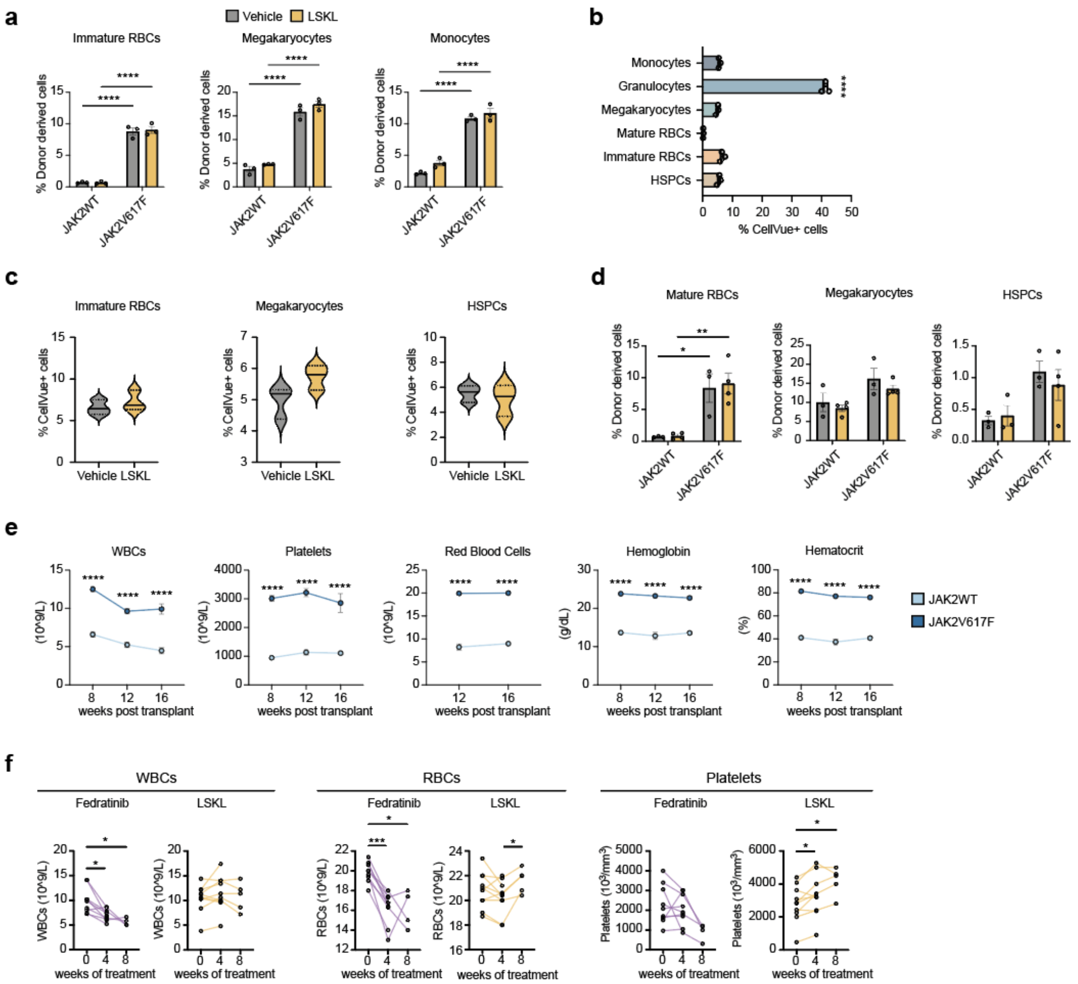


**Fig. S6 (related to Fig. 6): Pharmacological targeting of THBS1 synergizes with JAK inhibition to suppress myelofibrosis, halt fibrosis, and restore bone homeostasis.**

**a,** Flow cytometric analysis of donor-derived cells from BM organoids treated with vehicle or LSKL, showing frequencies of immature RBCs (CD45+CD235a+), megakaryocytes (CD45+CD61+) and monocytes (CD45+CD14+) derived from mutant and WT competitors. Two-way-ANOVA with post hoc Tukey's was used. **b,** Flow cytometric analysis of patient-derived CellVue-labelled primary CD34+ cells and their lineage bias toward granulocytes in the vehicle-treated BM organoids. One-way ANOVA with Dunnett's multiple comparison test was used. Significance shows the comparison of granulocytes versus every other cell population. **c,** Flow cytometric analysis of CellVue-labeled patient-derived cells following vehicle or LSKL treatment. **d,** Flow cytometric analysis of donor-derived cells from BM organoid-on-a-chip treated with vehicle or LSKL, showing frequencies of mature RBCs (CD45-CD235a+), megakaryocytes (CD45+CD61+) and HSPCs (CD45+CD34+) derived from mutant and WT competitors. Two-way-ANOVA with post hoc Tukey's was used. **e,** Blood counts over time showing white blood cells (WBC), red blood cells (RBCs), platelets, hemoglobin and hematocrit levels in both JAK2V617F and JAK2WT mice (n=5 mice per group). Two-way ANOVA followed by Sidak's multiple-comparisons test was used. **f,** Blood counts over time of treatment showing white blood cells (WBC), red blood cells (RBCs) and platelets. Each dot represents the value from one mouse. Two-way-ANOVA with post hoc Tukey's was used.
